# Metalloproteinase-dependent and TMPRSS2-independnt cell surface entry pathway of SARS-CoV-2 requires the furin-cleavage site and the S2 domain of spike protein

**DOI:** 10.1101/2021.12.14.472513

**Authors:** Mizuki Yamamoto, Jin Gohda, Ayako Kobayashi, Keiko Tomita, Youko Hirayama, Naohiko Koshikawa, Motoharu Seiki, Kentaro Semba, Tetsu Akiyama, Yasushi Kawaguchi, Jun-ichiro Inoue

## Abstract

The ongoing global vaccination program to prevent SARS-CoV-2 infection, the causative agent of COVID-19, has had significant success. However, recently virus variants have emerged that can evade the immunity in a host achieved through vaccination. Consequently, new therapeutic agents that can efficiently prevent infection from these new variants, and hence COVID-19 spread are urgently required. To achieve this, extensive characterization of virus-host cell interactions to identify effective therapeutic targets is warranted. Here, we report a cell surface entry pathway of SARS-CoV-2 that exists in a cell type-dependent manner is TMPRSS2-independent but sensitive to various broad-spectrum metalloproteinase inhibitors such as marimastat and prinomastat. Experiments with selective metalloproteinase inhibitors and gene-specific siRNAs revealed that a disintegrin and metalloproteinase 10 (ADAM10) is partially involved in the metalloproteinase pathway. Consistent with our finding that the pathway is unique to SARS-CoV-2 among highly pathogenic human coronaviruses, both the furin cleavage motif in the S1/S2 boundary and the S2 domain of SARS-CoV-2 spike protein are essential for metalloproteinase-dependent entry. In contrast, the two elements of SARS-CoV-2 independently contributed to TMPRSS2-dependent S2 priming. The metalloproteinase pathway is involved in SARS-CoV-2-induced syncytia formation and cytopathicity, leading us to theorize that it is also involved in the rapid spread of SARS-CoV-2 and the pathogenesis of COVID-19. Thus, targeting the metalloproteinase pathway in addition to the TMPRSS2 and endosome pathways could be an effective strategy by which to cure COVID-19 in the future.

**Author Summary:** To develop effective therapeutics against COVID-19, it is necessary to elucidate in detail the infection mechanism of the causative agent, SARS-CoV-2, including recently emerging variants. SARS-CoV-2 binds to the cell surface receptor ACE2 via the Spike protein, and then the Spike protein is cleaved by host proteases to enable entry. Selection of target cells by expression of these tissue-specific proteases contributes to pathogenesis. Here, we found that the metalloproteinase-mediated pathway is important for SARS-CoV-2 infection, variants included. This pathway requires both the prior cleavage of Spike into two domains and a specific sequence in the second domain S2, conditions met by SARS-CoV-2 but lacking in the related human coronavirus SARS-CoV. The contribution of several proteases, including metalloproteinases, to SARS-CoV-2 infection was cell type dependent, especially in cells derived from kidney, ovary, and endometrium, in which SARS-CoV-2 infection was metalloproteinase-dependent. In these cells, inhibition of metalloproteinases by treatment with marimastat or prinomastat, whose safety was previously confirmed in clinical trials, was important in preventing cell death. Our study provides new insights into the complex pathogenesis unique to COVID-19 and relevant to the development of effective therapies.

## Introduction

Severe acute respiratory syndrome coronavirus 2 (SARS-CoV-2), the causative agent of coronavirus disease 2019 (COVID-19), was first recognized in late 2019 and led to the development of a global pandemic in 2020[1]. Two other human coronaviruses, SARS-CoV[2, 3] and Middle East respiratory syndrome coronavirus (MERS-CoV)[4], are also capable of inducing lethal pneumonia and systemic symptoms. However, SARS-COV-2 has been found to also exhibit enhanced pathogenicity and transmissibility[5, 6]. Effective vaccines have been developed, and ongoing global vaccination programs have significantly curbed the spread of infection[7, 8]. However, current vaccinations may provide imperfect protection as new variants of the virus that can spread more easily and evade the host immunity achieved through vaccination have been reported[7, 9–11]. Furthermore, although several drugs that may provide effective treatments for COVID-19 are currently under clinical trial and awaiting approval[12, 13], it is currently unclear if daily life around the world will ever return to that of pre-COVID-19 times. Consequently, further extensive characterization of the virus and its interactions with host cells are required to develop vaccines and therapeutic agents that efficiently prevent infection from the new emerging highly infective variants so as to limit further worsening of COVID-19.

The initiation of SARS-CoV-2 entry requires two steps after its spike (S) protein is cleaved into S1 and S2 by furin-like proteases expressed in virus-producing cells prior to viral release[14–16]. First, the S protein binds to its receptor angiotensin converting enzyme 2 (ACE2) in the plasma membrane through its receptor-binding domain (RBD)[17, 18]. Second, the S2 protein is cleaved to generate S2′ by either cell surface transmembrane serine protease 2 (TMPRSS2)[19] or endosomal protease cathepsin-B/L[19, 20]. This cleavage is called priming, and exposes the fusion peptide within S2′, allowing it to stick into the plasma or endosomal membrane, resulting in fusion between the viral envelope and the cellular membrane (envelope fusion). This fusion allows viral RNA to enter the cytoplasm where it replicates. Whether SARS-CoV-2 viruses use the plasma membrane, the endosome pathway, or both is dependent on the cell type[19, 21, 22]. Furin-mediated cleavage at the S1/S2 boundary leads to efficient viral entry into airway cells[15, 16], where the TMPRSS2-dependent surface entry route dominates endosomal entry[19, 23].

In this study, we screened for inhibitors of SARS-CoV-2 infection and identified a cell surface entry pathway of SARS-CoV-2 that is TMPRSS2-independent but sensitive to various metalloproteinase inhibitors. Interestingly, the metalloproteinase-dependent pathway requires both the furin cleavage motif in the S1/S2 boundary and the S2 domain of SARS-CoV-2, which is unique to SARS-CoV-2. These results suggest that co-operation between furin and some metalloproteinases could be crucial for SARS-CoV-2 spread and disease development *in vivo*. Consequently, targeting the metalloproteinase-pathway in addition to the TMPRSS2 and cathepsin-B/L pathways could be an effective strategy to cure COVID-19.

## Results

### TMPRSS2-independent membrane fusion induced by the S protein of SARS-CoV-2 is blocked by metalloproteinase inhibitors

In this investigation, the screening system used to detect effective inhibitors of coronavirus infection included a quantitative cell fusion assay between effector cells expressing S protein and target cells expressing either ACE2 (for SARS-CoV and SARS-CoV-2)[24] or CD26 (for MERS-CoV)[25], with or without TMPRSS2 (S1a,b Fig). Quantitation was accomplished using the dual split chimeric reporter proteins (DSP)1-7 and DSP8-11, which contain both *Renilla* luciferase (RL) and green fluorescent protein (GFP) variants[26]. The DSP assay quantifies the degree of membrane fusion between the effector cells expressing DSP1-7 and the target cells expressing DSP8-11 based on the RL activity (S1c Fig). During analysis with the DSP cell fusion assay, a significant amount of ACE2-dependent but TMPRSS2-independent cell-cell fusion was induced by the S protein of SARS-CoV-2, but not by that of SARS- or MERS-CoV (Fig 1a,b). Consistent with this finding, the cell fusion with TMPRSS2 in the target cells induced by the S protein of SARS-CoV and MERS-CoV was completely blocked when TMPRSS2 was inhibited with 1 μM nafamostat, while approximately 20% of the fusion by the SARS-CoV-2 S protein remained, even in the presence of 10 μM nafamostat (Fig 1c). This amount of residual fusion was almost equal to that induced by the SARS-CoV-2 S protein in the absence of TMPRSS2 (Fig 1d).

**Fig 1.**
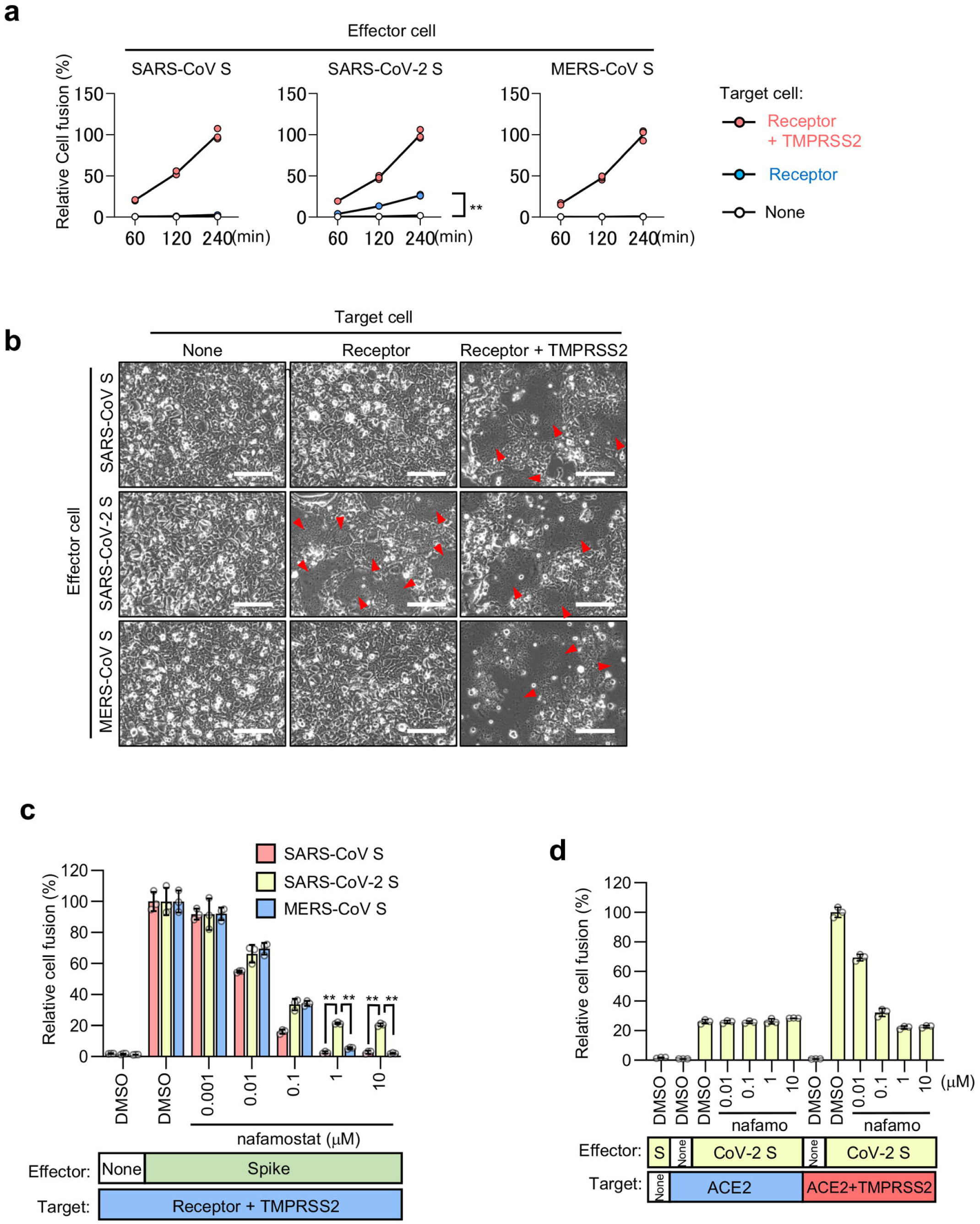
ACE2-dependent but TMPRSS2-independent membrane fusion activity of the SARS-CoV-2 S protein. **(a)** Cell fusion kinetics induced by the S proteins from SARS-CoV, SARS-CoV-2, and MERS-CoV were determined using the DSP assay. Target cells expressing ACE2 alone or together with TMPRSS2 were used for co-culturing with effector cells expressing SARS-CoV S and SARS-CoV-2 S, and cells expressing CD26 alone or together with TMPRSS2 were used for co-culturing with effector cells expressing MERS-CoV-S. Relative cell-fusion values were calculated by normalizing the RL activity of each co-culture to that of the co-culture with cells expressing both receptor and TMPRSS2 at 240 min, which was set to 100%. Values are means ± SD (*n* = 3/group). ** p < 0.01. (**b**) Phase contrast images of S protein-mediated cell fusion 16 h after co-culture. Red arrowheads indicate syncytia formation Scale bars, 100 μm. (**c**) Effect of nafamostat on the TMPRSS2-dependent cell fusion. Target cells expressing ACE2 with TMPRSS2 were used for co-culturing with effector cells expressing SARS-CoV S and SARS-CoV-2 S, and cells expressing CD26 with TMPRSS2 were used for co-culturing with effector cells expressing MERS-CoV-S. Relative cell-fusion values were calculated by normalizing the RL activity for each co-culture to that of the co-culture with cells expressing both receptor and TMPRSS2 in the presence of DMSO, which was set to 100%. Values are means ± SD (*n* = 3/group). ** p < 0.01. (**d**) Effects of the nafamostat on the TMPRSS2-independent or -dependent cell fusion. Target cells expressing ACE2 alone or together with TMPRSS2 were used for co-culturing with effector cells expressing SARS-CoV-2 S. Relative cell-fusion value was calculated by normalizing the RL activity for each co-culture to that of the co-culture with cells expressing both ACE2 and TMPRSS2 in the presence of DMSO, which was set to 100%. Values are means ± SD (*n* = 3/group). nafamo, nafamostat.

To explore the mechanism of TMPRSS2-independent membrane fusion, we screened the Validated Compound Library (1,630 clinically approved compounds and 1,885 pharmacologically active compounds) obtained from the Drug Discovery Initiative (The University of Tokyo). We aimed to identify compounds that preferentially inhibited SARS-CoV-2 S protein-induced TMPRSS2-independent fusion and not TMPRSS2- dependent fusion. We compared the relative fusion values of the target cells expressing both TMPRSS2 and ACE2 (X-axis of Fig 2a) with those of the target cells expressing ACE2 alone (Y-axis of Fig 2a), and chose for further validation compounds that limited the fusion without TMPRSS2 by less than 60% and allowed the fusion with TMPRSS2 by more than 70% (Fig 2a). The compounds selected included two metalloproteinase inhibitors (ilomastat, CTS-1027), three tyrosine kinase inhibitors (sunitinib, PD-166285, PD-173952), two checkpoint kinase inhibitors (PF-477736, AZD-7762), a protein kinase C inhibitor (midostaurin), and a hormonal contraceptive (algestone). Ilomastat and CTS-1027 preferentially inhibited the TMPRSS2-independent fusion in a dose-dependent manner without affecting TMPRSS2-dependent fusion (Fig 2b). Furthermore, the luciferase activities of the preformed DSP1-7/DSP8-11 complex were not affected, confirming the specificity of the DSP assay (S2a,b Fig). However, other compounds inhibited both the TMPRSS2-dependent and -independent fusions to similar degrees (S3 Fig). These data suggest that the metalloproteinase-dependent cell surface entry pathway (the metalloproteinase pathway) may be unique to SARS-CoV-2 among human pathogenic coronaviruses with high mortality rates. Considering that metalloproteinase inhibitors could thus possibly be used as prophylactic or therapeutic agents for COVID-19, we further demonstrated that marimastat[27] and prinomastat[28] (whose safety was previously confirmed in clinical trials to investigate their use as anticancer agents, such as CTS-1027[29]) can preferentially block the TMPRSS2-independent fusion induced by the SARS-CoV-2 S protein (Fig 2b and S2b Fig).

**Fig 2.**
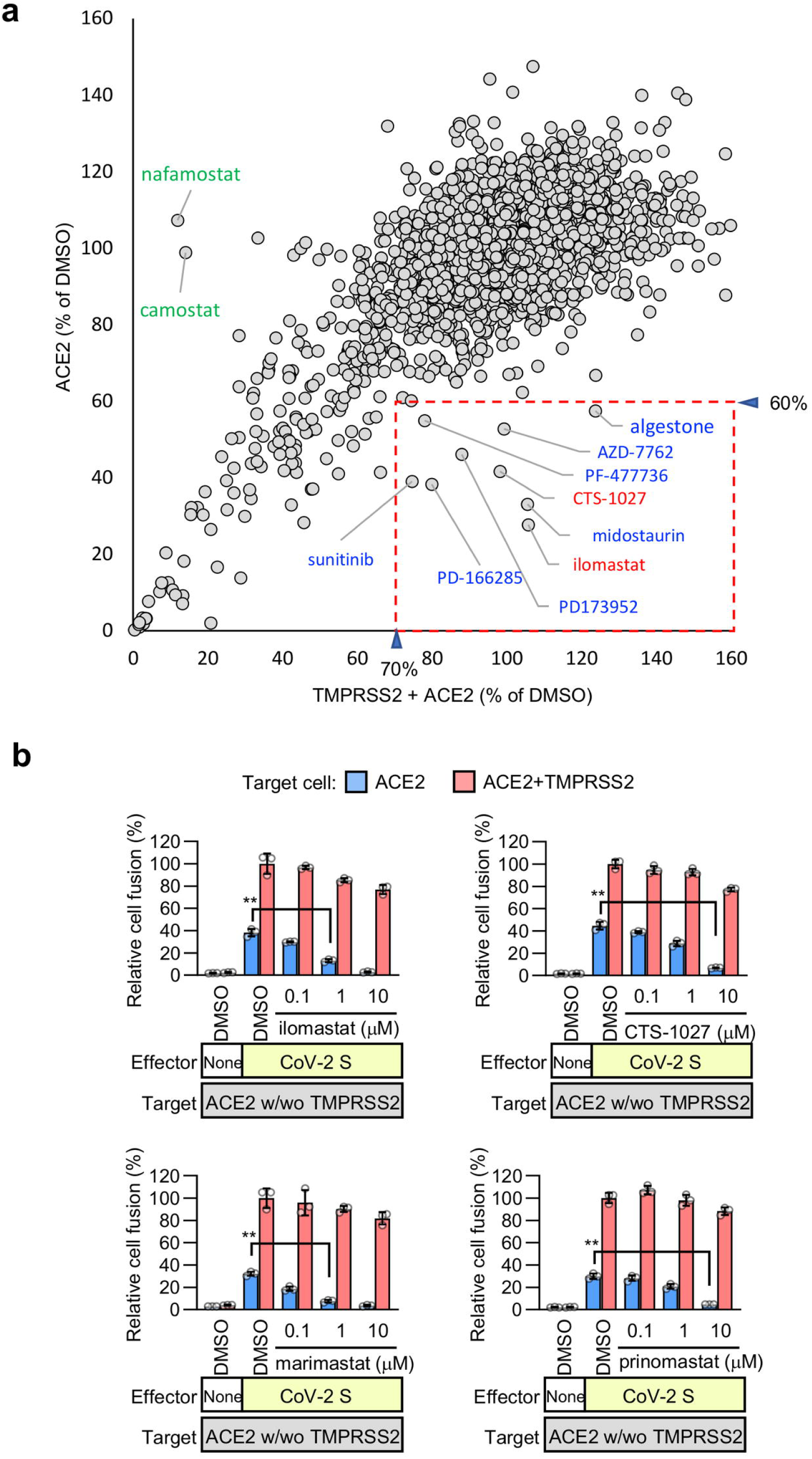
TMPRSS2-independent membrane fusion induced by the S protein of SARS-CoV-2 is blocked by various metalloproteinase inhibitors. **(a)** High-throughput screening of the Validated Compound Library (1,630 clinically approved compounds and 1,885 pharmacologically active compounds) in the DSP assay using the SARS-CoV-2 S protein. The x-axis shows the relative cell-fusion value using cells expressing both TMPRSS2 and ACE2 in the presence of each compound (1 µM in DMSO), n = 1. The y-axis shows the relative cell-fusion value using cells expressing ACE2 alone in the presence of each compound (1 µM in DMSO), n = 1. The relative cell-fusion value was calculated by normalizing the RL activity for each compound to that of the control assay (DMSO alone; set to 100%). Each dot represents an individual compound. Dots in the red-dashed box indicate compounds that preferentially inhibit TMPRSS2-independent membrane fusion. (< 30% inhibition of the relative cell fusion value using the target cells expressing both TMPRSS2 and ACE2 and > 40% inhibition of the relative cell-fusion value using the target cells expressing ACE2 alone. The compound names for the candidates are indicated. (**b**) Effects of the metalloproteinase inhibitors on cell fusion in the co-cultures of the cells expressing SARS-CoV-2 S protein with those expressing ACE2 alone or in combination with TMPRSS2. Relative cell-fusion values were calculated by normalizing the RL activity for each co-culture to that of the co-culture with cells expressing both ACE2 and TMPRSS2 in the presence of DMSO, which was set to 100%. Values are means ± SD (*n* = 3/group). ** p < 0.01.

### The metalloproteinase pathway is SARS-CoV-2 specific and cell type-dependent

We investigated whether the metalloproteinase pathway exists in SARS-CoV-2 S-bearing vesicular stomatitis virus (VSV) pseudovirus. The pseudovirus entry into the A704 cells (human kidney) was entirely blocked by 1 μM marimastat (Fig 3a). This indicates that the metalloproteinase pathway is involved in the entry of the virus, and that 1 μM marimastat could be used to determine if the pathway exists in other cells. Similarly, all entry pathways in OVISE cells (human ovary) were blocked by 25 μM E-64d (Fig 3a), indicating that it could be used to investigate the existence of cathepsin-B/L-dependent endosome pathways in other cells. Furthermore, the entry pathways in Calu-3 cells (human lung) were entirely blocked by 0.1 μM nafamostat (Fig 3a), which indicates that more than 0.1 μM nafamostat may be used to investigate the existence of the TMPRSS2-dependent surface entry pathways in other cells. Consequently, 1 μM marimastat, 25 μM E-64d, and 10 μM nafamostat were used to elucidate the patterns of the entry pathways in various cells. Marimastat significantly inhibited the pseudovirus entry into VeroE6 (African green monkey kidney), HEC50B (human endometrium), OVTOKO (human ovary), and A704 cells (Fig 3b). In addition to the metalloproteinase pathway, virus entry was partially inhibited by E-64d, and the combination of marimastat and E-64d showed additive effects in VeroE6, HEC50B, and OVTOKO cells (Fig 3b). These results suggest that the metalloproteinase and endosomal pathways are mutually independent. Neither marimastat nor nafamostat alone significantly inhibited the entry pathways of IGROV1 (human ovary), OUMS-23 (human colon), or OVISE, whereas E-64d significantly inhibited the entry pathways of IGROV1 and OUMS-23 cells and the overall entry pathway into the OVISE cells (Fig 3c). Interestingly, E-64d resistant entry in IGROV1 cells was inhibited by the combination of marimastat with E-64d while the E-64d resistant entry into OUMS-23 cells was inhibited by the combination of nafamostat and E-64d. These results indicate that the endosome entry pathway dominates these cells, while coexisting with either the metalloproteinase or TMPRSS2 surface pathway. Nafamostat inhibited the overall entry pathways into the Calu-3 and Caco-2 (human colon) cells, while marimastat and E-64d showed no inhibitory effects. (Fig 3d). Together, these findings show that the metalloproteinase-dependent cell surface entry pathway exists in a cell type-dependent manner and coexists with the endosome pathway in some cell lines. As we could not find cell lines with both metalloproteinase- and TMPRSS2-dependent cell surface entry pathways, we generated HEC50B cell lines ectopically expressing TMPRSS2 (HEC50B-TMPRSS2). In the HEC50B-TMPRSS2 cells, approximately 80% of the entry pathways were TMPRSS2-dependent, while the rest were predominantly metalloproteinase-dependent (Fig 3e). This indicates that the metalloproteinase pathway independently coexists with the TMPRSS2 dependent pathway. These results also suggest that there could be cells *in vivo* that naturally have both surface entry pathways.

**Fig 3.**
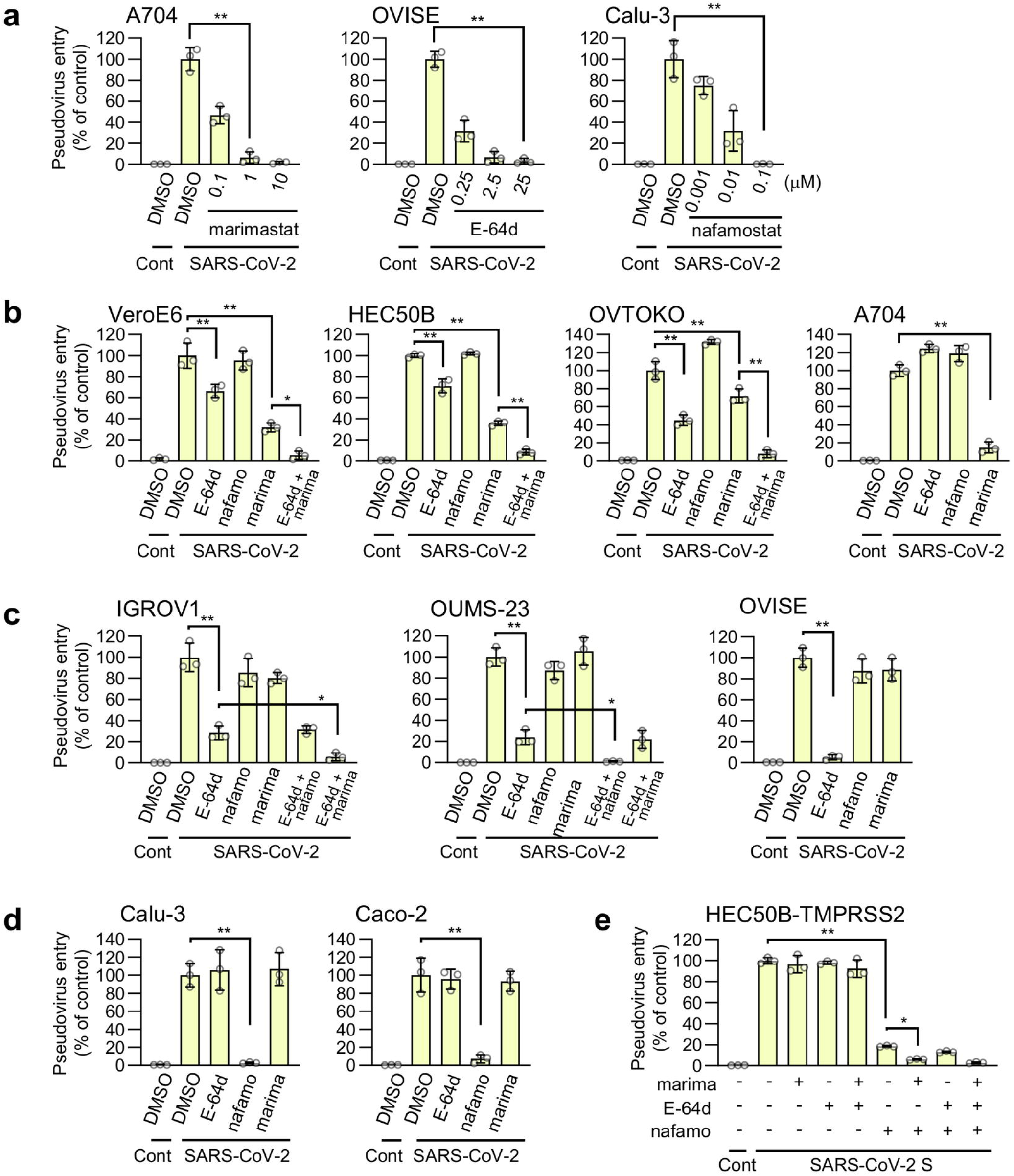
The metalloproteinase-dependent viral entry pathway is cell type-dependent. Effects of drugs on the entry of SARS-CoV-2 S-bearing vesicular stomatitis virus (VSV) pseudotype virus. The relative pseudovirus entry was calculated by normalizing the FL activity for each condition to the FL activity of cells infected with SARS-CoV-2 S-bearing pseudovirus in the presence of DMSO alone, which was set to 100%. Values are means ± SD (*n* = 3/group). * p < 0.05, ** p < 0.01. Cont: cells infected with pseudovirus without S protein; SARS-CoV-2: cells infected with SARS-CoV-2 S-bearing pseudovirus. E-64d: 25 μM E-64d, nafamo: 10 μM nafamostat, marima: 1 μM marimastat. (**a**) Effects of marimastat, E-64d, or nafamostat on the pseudovirus entry in A704, OVISE, and Calu-3 cells, respectively. (**b-e**) Effects of a single drug treatment or a combination treatment on the pseudovirus entry in VeroE6, HEC50B, OVTOKO and A704 cells (b), IGROV1, OUMS-23 and OVISE cells (c), Calu-3 and Caco-2 (d), and HEC50B-TMPRSS2 cells (e).

### The metalloproteinase pathway requires both the furin-cleavage site and S2 region of the SARS-CoV-2 S protein

Our results showed that metalloproteinase-dependent and TMPRSS2-independent cell-cell fusion was induced by the S protein of SARS-CoV-2 but not by that of SARS-CoV or MERS-CoV (Fig 1a,b). In line with these results, metalloproteinase-dependent entry was observed only when the pseudovirus bearing the S protein of SARS-CoV-2, but not SARS-CoV or MERS-CoV, was used in HEC50B (Fig 4a), A704 (S4a Fig), and VeroE6 cells (S4b Fig). Furthermore, pseudoviruses bearing the S protein of HCoV-NL63 and WIV1-CoV, which like SARS-CoV-2 use ACE2 as their receptor, cannot utilize the metalloproteinase pathway in HEC50B cells (Fig 4b). While SARS-CoV-2 S uses both the metalloproteinase and endosome pathways, SARS-CoV, MERS-CoV, HCoV-NL63, and WIV1-CoV S exclusively use the endosome pathway, and none of these S proteins can use the metalloproteinase or TMPRSS2 pathway in HEC50B (Fig 4a,b), A704 (S4a Fig), and VeroE6 cells (S4b Fig). The ability to use the metalloproteinase pathway and sensitivities against various protease inhibitors are conserved among the variants of SARS-CoV-2 we tested (S5 Fig).

**Fig 4.**
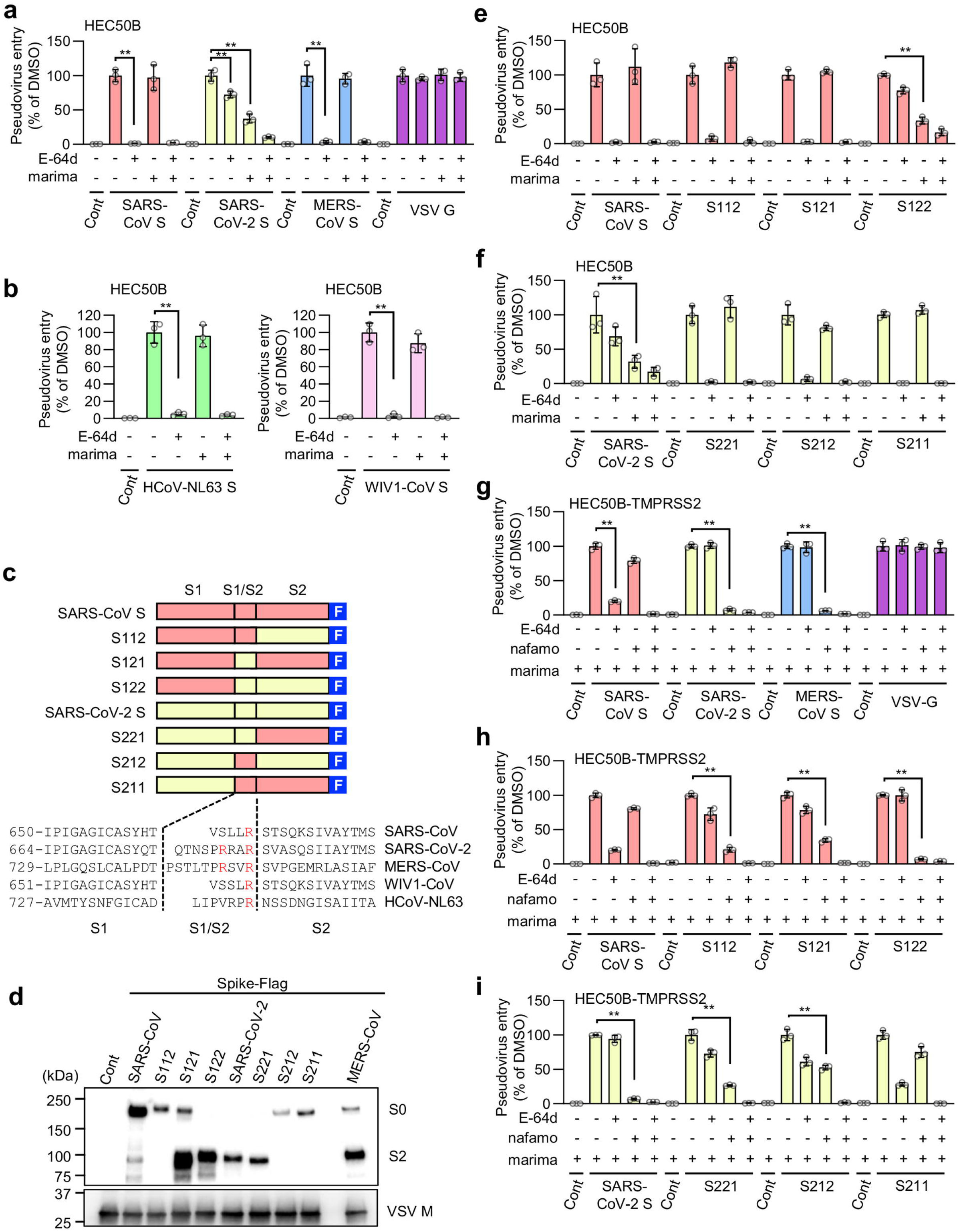
The metalloproteinase-dependent entry pathway requires both the furin-cleavage site and S2 region of the SARS-CoV-2 S protein. Effects of drugs on the entry of S protein-bearing vesicular stomatitis virus (VSV) pseudotype virus. The relative pseudovirus entry was calculated by normalizing the FL activity for each condition to the FL activity of cells infected with pseudovirus in the presence of DMSO alone, which was set to 100%. Values are means ± SD (*n* = 3/group). ** p < 0.01. Cont: cells infected with pseudovirus without S protein. E-64d: 25 μM E-64d, marima: 1 μM marimastat, nafamo: 10 μM nafamostat. (**a**) Effects of E-64d and marimastat on the entry of pseudoviruses bearing SARS-CoV S, SARS-CoV-2 S, MERS-CoV S, or VSV G in HEC50B cells. **b**, Effects of E-64d and marimastat on the entry of HCoV-NL63 S and WIV1-CoV S pseudovirus in HEC50B cells. **c**, Schematic illustration of C-terminally Flag-tagged chimeric S proteins in which the S1, S1/S2 boundary, and S2 domain from SARS-CoV S (red) or SARS-CoV-2 S (yellow) are indicated (top). Amino acid sequences of the residues around the S1/S2 boundary of the coronaviruses (bottom). Numbers refer to the amino acid residues. F: Flag-tag. Arginine residues in the S1/S2 cleavage site and furin cleavage motif are highlighted in red. (**d**) Expression of chimeric S protein in pseudoviruses. S proteins were detected using an anti-Flag-tag antibody that binds to a Flag-tag on the C-terminus of the S proteins (top). Detection of the vesicular stomatitis virus matrix protein (VSV M) served as the control (bottom). S0: uncleaved S protein; S2: cleaved S2 domain of the S protein. (**e, f**) Effects of E-64d and marimastat on the entry of pseudoviruses bearing chimeric S proteins in HEC50B cells. (**g**) Effects of E-64d and nafamostat on the entry of pseudoviruses bearing SARS-CoV S, SARS-CoV-2 S, MERS-CoV S, or VSV G in HEC50B-TMPRSS2 cells in the presence of marimastat. (**h, i**) Effects of E-64d and nafamostat on the entry of pseudoviruses bearing chimeric S proteins in HEC50B-TMPRSS2 cells in the presence of marimastat.

The S proteins of SARS-CoV-2 and MERS-CoV have furin cleavage sites (Arg-X-X-Arg) in their S1/S2 boundary area, and they were efficiently cleaved during the preparation of the pseudovirus (Fig 4c,d). In contrast, the S proteins of SARS-CoV, HCoV-NL63, and WIV1-CoV, which do not use the metalloproteinase pathway, do not have furin cleavage site and are not cleaved to any notable degree (Fig 4c,d and S6 Fig). Given that the MERS-CoV S protein does not use the metalloproteinase pathway (Fig 4a) even though it harbors a furin cleavage site and was efficiently cleaved, we speculated that furin-catalyzed S protein cleavage is a prerequisite but not sufficient for using the metalloproteinase pathway. To test this hypothesis, we generated pseudoviruses bearing chimeric S proteins in which the S1, S1/S2 boundary, and S2 domains were derived from either SARS-CoV or SARS-CoV-2 (Fig 4c,d). As expected, the S2 fragment of the C-terminal Flag-tagged S protein was mainly detected with anti-Flag antibody when pseudoviruses bearing S proteins with the furin-cleavage site (S121, S122, SARS-CoV-2 S (222), and S221) were analyzed (Fig 4d). In contrast, uncleaved S protein (S0) was mainly detected when S proteins without the furin-cleavage site (SARS-CoV S (111), S112, S212, and S211) were used (Fig 4d). When only section, the S1/S2 boundary or the S2 domain, was replaced with the corresponding domain of SARS-CoV-2 in the SARS-CoV S protein (S121, S112), the metalloproteinase pathway did not appear (Fig 4e). However, when both the S1/S2 and S2 domains were replaced with the corresponding domains of SARS-CoV-2 (S122), the metalloproteinase pathway appeared in addition to the endosome pathway (Fig 4e), similar to the pattern observed for bona fide SARS-CoV-2 S (S222) (Fig 4f). Furthermore, when either the S1/S2 or S2 domain was replaced with the corresponding domain of SARS-CoV in the SARS-CoV-2 S protein (S212, S221), the metalloproteinase pathway disappeared (Fig 4f). Similar requirements for the S1/S2 boundary and S2 domains for SARS-CoV-2 to use the metalloproteinase pathway were also observed in VeroE6 cells (S4c,d Fig). These results indicate that both the S1/S2 boundary and S2 domain of SARS-CoV-2 are strictly required for the virus to utilize the metalloproteinase pathway when the endosome pathway coexists.

To compare the structural requirements needed for the S protein to use the metalloproteinase pathway with those needed to use the TMPRSS2 pathway in cells with the endosome pathway as an alternative, we used HEC50B-TMPRSS2 and VeroE6 cells ectopically expressing TMPRSS2 (VeroE6-TMPRSS2). While both these cell types exhibit the TMPRSS2 and endosome pathways depending on the source of S protein (Fig 4g and S4e Fig), HEC50B-TMPRSS2 cells in addition maintained a significant amount of the metalloproteinase pathway (approximately 20% of the total entry pathway) (Fig 3e), whereas most of the metalloproteinase pathway disappeared in the VeroE6-TMPRSS2 cells when compared with their parental line (Fig 3b and S4e Fig). Given that the metalloproteinase pathway is dependent on the structural features of the S protein, its blockade by marimastat will promote protein structural requirements for S protein to use the TMPRSS2 pathway in HEC50B-TMPRSS2 (Fig 4g-i) but not in VeroE6-TMPRSS2 cells (S4e-g Fig). In both cell types, SARS-CoV-2 predominantly used the TMPRSS2 entry pathway, while SARS-CoV used the endosome pathway (Fig 4g and S4e Fig). However, nafamostat partially, but more efficiently, inhibited the entry of S112 and S121 pseudoviruses when compared with SARS-CoV (S111), while it inhibited S122 virus entry almost completely (Fig 4h and S4f Fig). Furthermore, when compared with SARS-CoV-2 (S222), nafamostat only partially inhibited S221 or S212 virus entry, while it scarcely inhibited S211 virus entry (Fig 4i and S4g Fig). These results indicate that the S1/S2 boundary and S2 domain of SARS-CoV-2 additively contribute to the ability of the virus to use the TMPRSS2 pathway. Together, these results show that although both the TMPRSS2 and the metalloproteinase pathways undergo priming of the S protein at the cell surface, the structural requirements of the S protein for efficient priming differ between the metalloproteinase and TMPRSS2 pathways.

### Possible involvement of ADAM-10 in the metalloproteinase-dependent entry of SARS-CoV-2

In addition to marimastat, other metalloproteinase inhibitors, including prinomastat, ilomastat, and CTS-1027, which block the TMPRSS2-independent cell-cell fusion induced by SARS-CoV-2 S (Fig 2b), inhibited the metalloproteinase-dependent entry of SARS-CoV-2 pseudovirus in VeroE6, HEC50B, and A704 cells (Fig 5a). Since these inhibitors exhibited broad specificity[30–33], selective inhibitors were then used to narrow down the metalloproteinases involved in the metalloproteinase-dependent entry pathway. The VeroE6 and HEC50B cells had significantly E-64d sensitive endosome pathways but the A704 cells did not (Fig 3b). Consequently, selective metalloproteinase inhibitors were tested in the presence of E-64d so that the reduction in the metalloproteinase pathway could be easily recognized in VeroE6 and HEC50B cells (Fig 5a). Similar inhibitory patterns were observed in all three cell lines tested (Fig 5a), and their viabilities were not affected by any of the metalloproteinase inhibitors at the concentrations used in the experiment (S7a-d Fig). These results suggest that the metalloproteinases involved in the pathway are likely to be common to all three cell lines. GW280264X[34] (ADAM10/17 inhibitor) and GI1254023X[34, 35] (MMP9/ADAM10 inhibitor) significantly inhibited the metalloproteinase pathway, whereas TAPI2[36] (ADAM-17 inhibitor) and BK-1361[37] (ADAM8 inhibitor) did not (Fig 5a). This suggests that ADAM10 may be involved in the ADAM family. MMP408[30] (MMP3/12/13 inhibitor) and MMP2/9 inhibitor I[38] scarcely affected virus entry, whereas UK370106[39] (MMP3/12 inhibitor) and MMP9 inhibitor I[40] were significantly inhibitory (Fig5a), suggesting that MMP3/9/12/13 may not be crucial, but that the unidentified metalloproteinase, which could be inhibited by UK370106 or MMP9 inhibitor I, may be involved in the pathway in cooperation with ADAM10. MLN-4760[41] (ACE2 inhibitor) did not inhibit virus entry (Fig 5a), indicating that the catalytic activity of ACE2, to which the S protein directly binds as a receptor, is not involved. To further confirm the involvement of ADAM10 in the metalloproteinase-dependent pathway, ADAM10 was depleted by siRNA in HEC50B cells. Three independent siRNAs effectively suppressed the expression of both the precursor and active forms of ADAM10 (Fig 5b). An ADAM10 knockdown significantly inhibited SARS-CoV-2 pseudovirus entry, while the entry of SARS-CoV, MERS-CoV, and VSVG pseudoviruses were not affected (Fig 5c), indicating that ADAM10 plays a role unique to SARS-CoV-2 in viral entry. Furthermore, we examined the effects of the ADAM10 knockdown on the entry pathway patterns by treating siRNA-transfected cells with either E-64d, marimastat, or a combination of both. The combination treatment with E-64d and marimastat led to an additive effect for the single treatments, resulting in the complete inhibition of viral entry in both cells with normal ADAM10 expression and those with reduced ADAM10 expression (Fig 5d). These results indicated that E-64d-resistant viral entry is a metalloproteinase-dependent pathway, while marimastat-resistant viral entry is dependent on the endosome pathway. The ADAM10-knockdown significantly inhibited the metalloproteinase pathway (Fig 5e, E64-d treatment) while the ADAM10-knockdown had only a modest effect on the endosome pathway (Fig 5e, marimastat-treatment), indicating that ADAM10 is involved in the metalloproteinase-dependent entry pathway of SARS-CoV-2.

**Fig 5.**
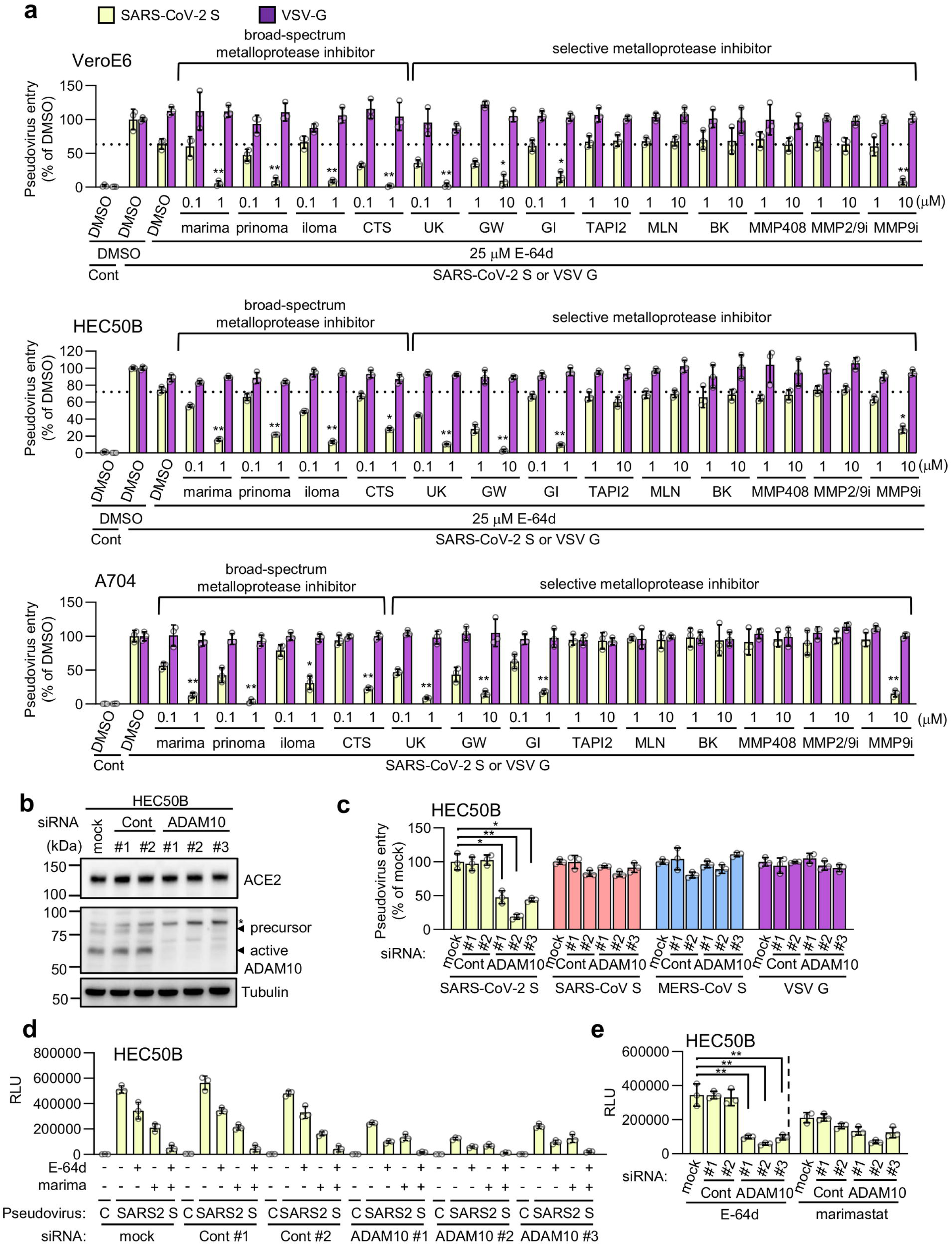
Possible involvement of ADAM-10 in the metalloproteinase-dependent entry of SARS-CoV-2. **(a)** Effects of metalloprotease inhibitors on the entry of pseudoviruses bearing SARS-CoV-2 S or VSV G in VeroE6 and HEC50B cells in the presence of E-64d, and A704 cells in the absence of E-64d. The relative pseudovirus entry was calculated by normalizing the FL activity for each condition to the FL activity of cells infected with pseudovirus in the presence of DMSO alone, which was set to 100%. Values are means ± SD (*n* = 3/group). Data were compared with those obtained from cells infected pseudoviruses bearing SARS-CoV-2 S in the presence of E-64d for HEC50B and VeroE6, and in the presence of DMSO alone for A704. * p < 0.05, ** p < 0.01. Cont: cells infected with pseudovirus without S protein. marima: marimastat, prinoma: prinomastat, iloma: ilomastat, CTS: CTS-1027, UK: UK370106, GW: GW280264X, GI: GI254023X, MLN: MLN-4760, BK: BK-1361, MMP2/9i: MMP2/9 inhibitor I, MMP9i: MMP9 inhibitor I. **(b)** Effects of the ADAM10 knockdown on ACE2 (top), ADAM10 (middle), and tubulin (bottom) expression. HEC50B cells were transfected with two distinct control siRNAs or three distinct siRNAs against Adam10 for 48 h. (**c**) The effect of the ADAM10 knockdown on the entry of pseudoviruses bearing SARS-CoV-2 S, SARS-CoV S, MERS-CoV S, or VSV G. HEC50B cells were transfected with siRNAs for 48 h and then infected with pseudoviruses. The relative pseudovirus entry was calculated by normalizing the FL activity for each condition to the FL activity of cells infected with pseudovirus in the absence of siRNA (mock), which was set to 100%. Values are means ± SD (*n* = 3/group). * p < 0.05, ** p < 0.01. (**d, e**) Effect of ADAM10 knockdown on the patterns of the entry pathways for SARS-CoV-2 S pseudovirus in HEC50B cells. HEC50B cells were transfected with siRNAs for 48 h and then infected with pseudoviruses in the presence of drugs. Values are means ± SD (*n* = 3/group). ** p < 0.01. E-64d: 25 μM E-64d, marima: 1 μM marimastat. Data are displayed as the conditions of siRNA treatment (d) and drug treatment (e).

Recently, it was reported that ACE2 shedding by ADAM17 promotes SARS-CoV-2 infection[42]. While a CRISPR/Cas9-mediated knockout of ADAM17 enhanced the accumulation of cellular ACE2 in HEC50B cells due to the inhibition of ACE2 shedding (S8a Fig), SARS-CoV-2 S pseudovirus entry was increased, and this was probably because of the enhanced binding of virus to the cell surface ACE2 (S8b Fig). However, the patterns of the metalloproteinase and endosome pathways were similar between the wild-type and ADAM17 knockout cells (S8c Fig), suggesting that ADAM17 may not be involved in metalloproteinase-dependent virus entry in HEC50B cells.

### The metalloproteinase-dependent entry pathway of authentic SARS-CoV-2 is involved in syncytia formation and cytopathicity

To confirm the involvement of the metalloproteinase pathway in authentic SARS-CoV-2 entry, we first evaluated the effects of marimastat and prinomastat on the amount of cytoplasmic viral RNA transcribed from the N gene after infection. Both inhibitors significantly suppressed SARS-CoV-2 infection (Fig 6a). The IC_50_ values of the marimastat and prinomastat were 160 nM and 130 nM in HEC50B cells, and 150 nM and 250 nM in the A704 cells, respectively. The IC_50_ value of the marimastat in the VeroE6 cells was 340 nM. Consistent with the results from the pseudoviruses experiments, nafamostat showed a marked inhibitory effect on the Calu-3 cells but not on the HEC50B, A704, or VeroE6 cells (S9a Fig). In contrast, 25 μM E-64d and 10 mM NH_4_Cl, which inhibits endosome-lysosome system acidification[43], significantly suppressed SARS-CoV-2 infection in the HEC50B, A704, and VeroE6 cells (S9b,c Fig). This indicated that the endosome pathway coexists with the metalloproteinase pathway to contribute to authentic SARS-CoV-2 infection in these cells. Combination treatments with E-64d/marimastat or NH_4_Cl/marimastat showed much stronger inhibitory effects than the treatment with each drug alone (Fig 6b). Similarly, combination treatments with nafamostat and marimastat or nafamostat and E-64d showed stronger inhibitory effects than the nafamostat treatment alone in the HEC50B-TMPRSS2 cells (Fig 6c). Furthermore, when all three drugs were combined, they had a much stronger inhibitory effect on viral infection when compared with the two-drug combinations (Fig 6c). These results strongly suggest that drugs that block the metalloproteinase pathway are effective for COVID-19 treatment.

**Fig 6.**
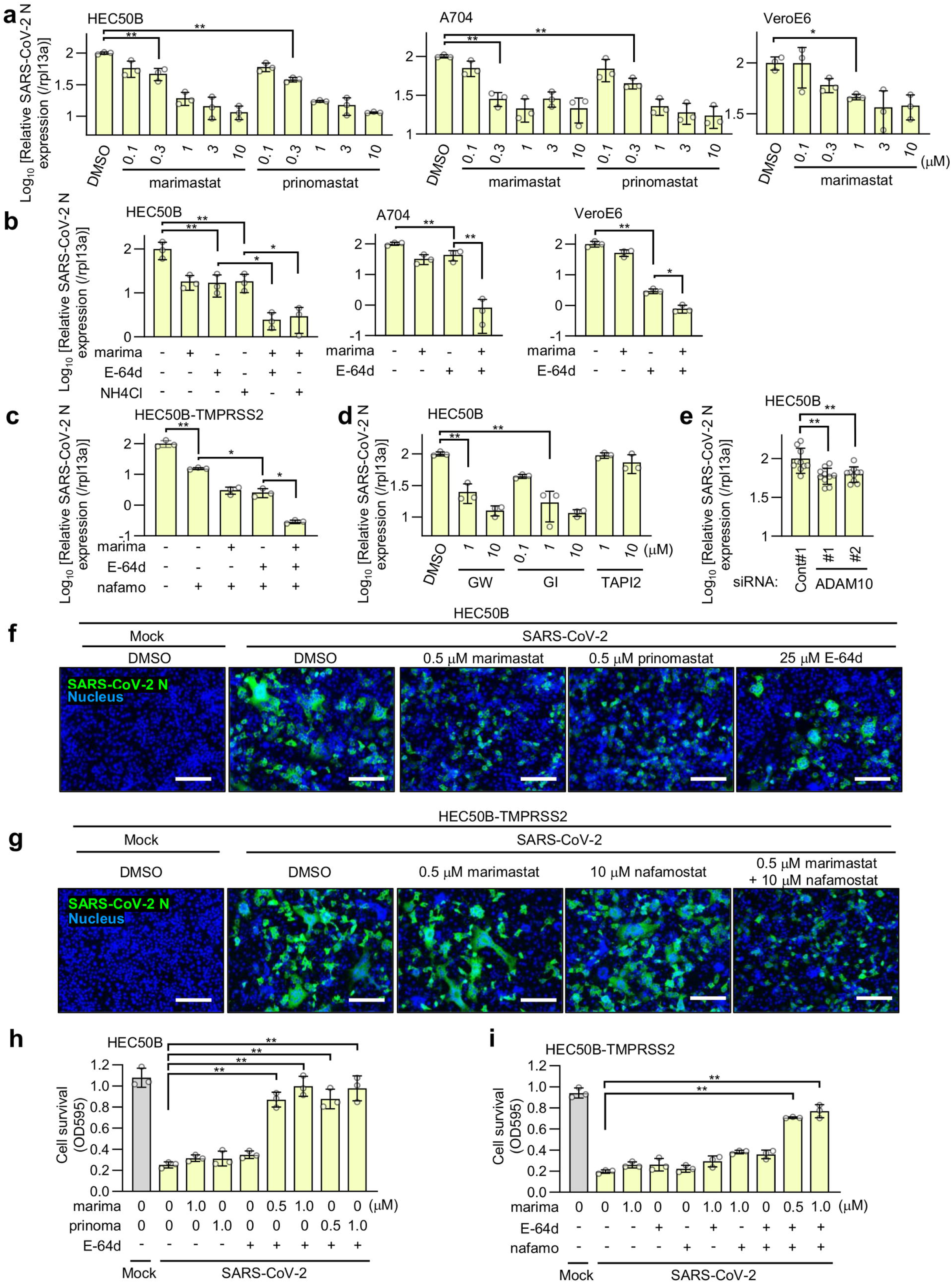
The metalloproteinase-dependent entry pathway of authentic SARS-CoV-2 is involved in syncytia formation and cytopathicity. Effects of the drugs on the cytoplasmic viral RNA after SARS-CoV-2 infection. The relative amount of viral RNA in the cells was normalized to cellular *Rpl13a* mRNA expression. Values are means ± SD (*n* = 3/group in a-d, *n* = 10/group in e). * p < 0.05, ** p < 0.01. (**a**) Effects of marimastat or prinomastat on SARS-CoV-2 infection in HEC50B, A704, and VeroE6 cells. (**b**) Effects of marimastat and the inhibitor of the endosome pathway on SARS-CoV-2 infection in HEC50B, A704, and VeroE6 cells. marima: 1 μM marimastat, E-64d: 25 μM E-64d, NH4Cl: 10 mM NH4Cl. (**c**) Effect of marimastat, E-64d, and nafamostat on SARS-CoV-2 infection in HEC50B-TMPRSS2 cells. marima: 1 μM marimastat, E-64d: 25 μM E-64d, nafamo: 10 μM nafamostat. (**d**) Effects of selective metalloprotease inhibitors on SARS-CoV-2 infection in HEC50B cells. GW: GW280264X, GI: GI254023X. (**e**) Effect of ADAM10 knockdown on SARS-CoV-2 infection in HEC50B cells. (**f, g**) Effects of drugs on SARS-CoV-2-induced syncytia formation in HEC50B (f) and HEC50B-TMPRSS2 (g) cells. Cells were stained with anti-SARS-CoV-2 N antibody (green) 24 h after infection. Nuclei were stained with Hoechst 33342 (blue). Scale bars, 200 μm. (**h, i**) Effects of drugs on SARS-CoV-2-induced cytopathicity in HEC50B (h) and HEC50B-TMPRSS2 (i) cells. marima: marimastat, prinoma: prinomastat. E-64d: 25 μM E-64d, nafamo: 10 μM nafamostat. Values are means ± SD (*n* = 3/group). ** p < 0.01.

Next, we examined whether ADAM10 is involved in SARS-CoV-2 infection. GW280264X[34] (ADAM10/17 inhibitor) and GI1254023X[34, 35] (MMP9/ADAM10 inhibitor) significantly suppressed SARS-CoV-2 infection, whereas TAPI2[36] (ADAM-17 inhibitor) did not (Fig 6d). Moreover, the ADAM10 knockdown by siRNA suppressed SARS-CoV-2 infection by approximately 40% (Fig 6e), indicating that ADAM10 is partially involved. This effect was smaller than that for the various metalloproteinase inhibitors (Fig 6a,d), suggesting that metalloproteinases other than ADAM10 are also involved in this pathway.

The ability of SARS-CoV-2 to form syncytia and induce cytopathicity is thought to be related to its pathogenesis[44, 45]. To determine whether the metalloproteinase-dependent pathway is involved in syncytia formation, we first used HEC50B cells as a representative for cells that predominantly use the metalloproteinase and endosome pathways. Interestingly, the SARS-CoV-2-induced syncytia formation in HEC50B cells 24 h after infection was significantly blocked by 500 nM of marimastat and prinomastat but not notably affected by 25 μM E-64d (Fig 6f and S10 Fig). These results indicate that the metalloproteinase-dependent pathway, but not the endosome pathway, is crucial for syncytium formation, although both pathways similarly reduce viral infection (Fig 6b). Given that the metalloproteinases are normally localized at the cell surface, we used HEC50B-TMPRSS2 cells, which have cell surface TMPRSS2 and metalloproteinase pathways, to investigate their involvement in syncytia formation when they coexist. The SARS-CoV-2-induced syncytia formation in the HEC50B-TMPRSS2 cells was not significantly inhibited by marimastat or nafamostat alone, but was clearly inhibited by the combined treatment (Fig 6g). These results suggest that the metalloproteinase and TMPRSS2 pathways cooperate to form syncytia. Next, we addressed the role of the metalloproteinase pathway in SARS-CoV-2-induced cytotoxicity. SARS-CoV-2-induced cytopathicity of HEC50B cells 3 d after infection was not inhibited by either E-64d, marimastat, or prinomastat alone, but was significantly blocked when cells were treated with E-64d in combination with either marimastat or prinomastat (Fig 6h). In addition, SARS-CoV-2-induced cytopathicity of the HEC50B-TMPRSS2 cells was not inhibited by either E-64d, nafamostat, or marimastat alone, but was significantly blocked when cells were treated with a combination of all three drugs (Fig 6i). These results strongly suggest that the inhibition of the metalloproteinase pathway is crucial to block syncytia formation and cytopathicity *in vivo*, and consequently, that the metalloproteinase pathway is likely to be involved in the pathogenesis of COVID-19.

## Discussion

Previous studies of the S proteins found in SARS-CoV-2, SARS-CoV, and MERS-CoV have shown that the priming of the S2 domain, catalyzed by either TMPRSS2 on the plasma membrane or cathepsin-B/L in endosomes, results in the fusion peptide protruding to induce envelope fusion, thereby establishing viral entry through the plasma membrane or endosomal membrane, respectively[14, 19, 20]. In this study, we have demonstrated that SARS-CoV-2, unlike SARS-CoV or MERS-CoV, has a unique TMPRSS2-independent cell surface entry pathway, which is sensitive to various metalloproteinase inhibitors including ilomastat, CTS-1027, marimastat, and prinomastat, but resistant to previously known inhibitors of SARS-CoV-2 entry, such as nafamostat[24, 46] and E-64d[19]. As a representative of these broad-spectrum metalloproteinase inhibitors, we chose marimastat to investigate the cell type-dependent distribution of the metalloproteinase pathway by measuring pseudovirus infection. A significant proportion of the entry pathway is metalloproteinase-dependent in A704 (kidney), HEC50B (endometrium), OVTOKO (ovary), and VeroE6 (kidney) cells. Only a small proportion was metalloproteinase-dependent in IGROV1 (ovary) cells, while the metalloproteinase pathway was not detected in OMUS-23 (colon), OVISE (ovary), Calu-3 (lung), and Caco-2 (colon) cells. These results indicate that the metalloproteinase pathway of SARS-CoV-2 is cell-type specific and independently coexists with other entry pathways, including the TMPRSS2-dependent surface pathway and the endosome pathway. The kidney[47, 48], ovary[49, 50], and endometrium[51] are known to express ACE2. Furthermore, SARS-CoV-2 can infect the kidney[48, 52] and induce acute kidney injury[53] in COVID-19 patients. Although SARS-CoV-2 infection of the ovary or endometrium has not previously been reported, the metalloproteinase-dependent infection pathway may contribute to the pathogenesis of COVID-19, especially multiple organ failure. The metalloproteinase pathway is thus a potential target for future COVID-19 therapies.

The S1/S2 boundary of SARS-CoV-2 contains the furin cleavage motif (Arg-X-X-Arg), while that of SARS-CoV contains only a single Arg. It has been reported that the motif greatly increases the efficiency of S1/S2 cleavage[15, 16], leading to enhanced viral transmission both *in vitro*[15, 16, 23] and *in vivo*[54, 55]. This may be partially due to the enhanced availability of S2 to TMPRSS2, due to the dissociation of S1[56, 57]. We have shown that the furin cleavage motif is required for the metalloproteinase pathway, and we propose that the induction of metalloproteinase-induced S2 priming is another role of furin-mediated S1/S2 cleavage in enhanced viral transmission. Therefore, the metalloproteinase-dependent entry pathway, which is unique among highly pathogenic coronaviruses, is likely to be associated with the rapid spread of SARS-CoV-2. Interestingly, experiments using pseudoviruses bearing chimeric S proteins between SARS-CoV (without the metalloproteinase pathway) and SARS-CoV-2 (with the metalloproteinase pathway) revealed that both the S1/S2 boundary of SARS-CoV-2 and the S2 domain of SARS-CoV-2 S are essential for metalloproteinase-dependent entry. In contrast, the two domains of SARS-CoV-2 independently contributed to TMPRSS2-dependent S2 priming. This discrepancy may be partially due to the difference in the substrate recognition properties of the priming proteases in the two pathways. The S112 pseudovirus (a VSV pseudovirus bearing SARS-CoV S mutant, in which the S2 region was replaced with the corresponding domain of SARS-CoV-2) can use the TMPRSS2 pathway more efficiently than the SARS-CoV S pseudovirus, which suggests that TMPRSS2 may be partially accessible to the priming site in SARS-CoV-2 (C-terminal of Arg815) but not to that in SARS-CoV (C-terminal of Arg797) without S1 dissociation. In contrast, the putative priming protease in the metalloproteinase pathway, which may not be a metalloproteinase but a protease activated by metalloproteinases, can access the priming site only when the site occurs within the contextual characteristics of SARS-CoV-2 S2, and S1/S2 is cleaved to allow S1 dissociation. Determination of the priming site in the metalloproteinase pathway and identification of the critical amino acid residues generating the structural characteristics of SARS-CoV-2 S2 that allow metalloproteinase-dependent priming are required to understand its molecular mechanisms for the two distinct surface entry pathways. From an evolutionary perspective, SARS-CoV-2 acquired the metalloproteinase pathway by introducing mutations into the S2 region, which may have contributed to the SARS-CoV-2 pandemic. The function of point mutations in the S2 domain have not yet been fully analyzed in comparison to those in the S1 domain, which cause escape from neutralizing antibodies[58]. However, various point mutations in the S2 domain may play important roles in increasing the efficiency of infection and disease progression and the generation of highly infectious variants.

Using selective metalloproteinase inhibitors and ADAM10 knockdowns generated using specific siRNAs, we have demonstrated that ADAM10 plays an important role in the metalloproteinase pathway. ADAM10 is ubiquitously expressed in various tissues[59] and cell lines[60], and functionally regulates cell differentiation and proliferation by cleaving ligands and receptors such as epidermal growth factor (EGF), heparin-binding EGF-like growth factor (HB-EGF), and Notch[61]. ADAM10 is thus likely to contribute to SARS-CoV-2 infection in various organs. ADAM17, similar to ADAM10, is also known as a metalloproteinase that induces the cleavage of receptors and ligands, and both can cleave common substrates such as Notch and HB-EGF[61]. It has been reported that ACE2 shedding by ADAM17 promotes SARS-CoV-2 infection[42]. Although we observed that ADAM17 depletion resulted in the cellular accumulation of ACE2, which is indicative of reduced ACE2 shedding, total SARS-CoV-2 pseudovirus infection was unexpectedly augmented and the relative contribution of the metalloproteinase pathway was not affected. ADAM10 and ADAM17 thus both play crucial but distinct roles in SARS-CoV-2 infection. It has recently been reported that ADAM9 inhibition decreases SARS-CoV-2 infection *in vitro*[62]. Although the involvement of ADAM9 in viral entry is not clear, the results suggest that a group of metalloproteinases cooperate in the metalloproteinase pathway. This may indicate that the observed ADAM10 depletion-induced inhibition was a part of the maximum inhibition by various metalloproteinase inhibitors. A recent report also showed that MMP12 knockouts inhibited SARS-CoV-2 infection *in vitro*[63]. However, MMP408[30], an inhibitor of MMP12, did not prevent SARS-CoV-2 infection in various cell lines in this investigation, suggesting that metalloproteinases involved in the metalloproteinase pathway may differ in a cell type-dependent manner. Further studies are required to identify the functional metalloproteinases that are involved in the metalloproteinase pathway.

Various compounds are reported to inhibit SARS-CoV-2 infection by inhibiting envelope fusion *in vitro*. However, camostat[64], an inhibitor of the TMPRSS2- dependent surface entry, and hydroxychloroquine[65, 66], an inhibitor of the endosomal pathway, have failed to show sufficient therapeutic efficacy in clinical trials. Our entry pathway analysis revealed that the metalloproteinase surface pathway coexists with the TMPRSS2 pathway and/or the endosome pathway in various cell types. Furthermore, in HEC50B and HEC50B-TMPRSS2 cells, cell death could not be inhibited unless all entry pathways in each cell were inhibited using inhibitor co-treatments for each pathway. Therefore, future clinical trials on virus entry in which the TMPRSS2, metalloproteinase, and endosome pathways are all efficiently blocked, need to be conducted. We propose that to address this challenge both marimastat and prinomastat should be utilized in clinical trials. The mean maximum plasma concentration (C_max_) at a reasonably well-tolerated dose was 590 nM for marimastat[27] and 680 nM for prinomastat[28]. Furthermore, we demonstrated that these two drugs significantly inhibited SARS-CoV-2 infection at concentrations lower than their C_max_ values. These metalloprotease inhibitors, in combination with other protease inhibitors targeting the TMPRSS2 and endosome pathways, may effectively inhibit SARS-CoV-2 infection in various tissues and cure COVID-19. Recently, there has been concern about the spread of SARS-CoV-2 variants, such as the delta strain, as they reduce the effectiveness of the neutralizing antibodies produced by vaccination[7, 9, 10]. Since sensitivities against marimastat, nafamostat, and E-64d have been conserved in the various variants investigated so far, the strategies to use protease inhibitors for COVID-19 treatment are likely to be significantly effective against these variants. The results of this study may contribute to the development of COVID-19 treatments targeting viral entry pathways.

## Materials and Methods

### Cell lines, viruses, and reagents

VeroE6 (CRL-1586), 293T (CRL-3216), A704 (HTB-45) and Calu-3 (HTB-55) cells were obtained from the American Type Culture Collection (Rockville, MD, USA). OVTOKO (JCRB1048), OVISE (JCRB1043), HEC50B (JCRB1145), VeroE6-TMPRSS2 (JCRB1819)[67], and OUMS-23 (JCRB1022) cells were obtained from the Japanese Collection of Research Bioresources Cell Bank (Osaka, Japan). IGROV1 cells (SCC203) were purchased from Merck (Darmstadt, Germany) and Caco-2 cells (RCB0988) were obtained from the RIKEN BioResource Research Center (Tsukuba, Japan). A704, Calu-3, VeroE6, HEC50B, and Caco-2 cells were maintained in Eagle’s minimum essential medium (EMEM; 055-08975, FUJIFILM Wako Pure Chemical, Osaka, Japan) containing 15% fetal bovine serum (FBS). OUMS-23, IGROV1, and 293T cells were maintained in Dulbecco’s modified Eagle’s medium (DMEM; 041-30081, FUJIFILM Wako Pure Chemical) containing 10% FBS. VeroE6-TMPRSS2 (JCRB1819) cells were cultured in DMEM containing 10% FBS and 1 mg/mL G418. OVISE and OVTOKO cells were maintained in Roswell Park Memorial Institute (RPMI)-1640 medium (189-02025, FUJIFILM Wako Pure Chemical) containing 10% FBS. A pair of previously described 293FT-based reporter cell lines that stably express individual split reporters (DSP1-7 and DSP8-11 proteins)[68] were maintained in DMEM containing 10% FBS and 1 μg/mL puromycin. To establish stable cell lines expressing the S protein of SARS-CoV, SARS-CoV-2, or MERS-CoV, recombinant pseudotype lentiviruses were produced in 293T cells with psPAX2 packaging plasmid, vesicular stomatitis virus (VSV)-G-expressing plasmid and lentiviral transfer plasmid expressing S protein. To establish stable cell lines expressing ACE2 or CD26 with TMPRSS2, recombinant pseudotype lentiviruses expressing one of the proteins were produced using 293T cells with psPAX2 packaging plasmid and VSV-G-expressing plasmid. The 293FT-derived reporter cells infected with the pseudotype viruses were selected with 1 μg/mL puromycin, 10 μg/mL blasticidin, and 300 μg/mL hygromycin for at least 1 week. These bulk-selected cells were used for fusion assays. To establish HEC50B cells expressing TMPRSS2 (HEC50B-TMPRSS2), recombinant pseudotype lentivirus expressing TMPRSS2 was produced using 293T cells with psPAX2 packaging plasmid and VSV-G-expressing plasmid. HEC50B cells infected with pseudotype viruses were selected with 300 μg/mL hygromycin for at least 1 week. The SARS-CoV-2 isolate (UT-NCGM02/Human/2020/Tokyo)[69] was propagated in VeroE6-TMPRSS2 (JCRB1819) cells in DMEM containing 5% FBS. Titers were determined with plaque assays using VeroE6/TMPRSS2 (JCRB1819) cells. Negative control No.1 siRNA (4390843), negative control No.2 siRNA (4390846), and three distanced ADAM10-specific siRNAs were purchased from Thermo Fisher Scientific (MA, USA) The siRNA sequences used were 5′-UCA CCU UGU UCU ACC AUU CCA (S1004, ADAM10#1); 5′-UAA CCU CUA AAA UCG UUG CAA (S1005, ADAM10#2); and 5′-UAC GGA UUC CGG AGA AGU CTG (S1006, ADAM10#3) for the ADAM10 knockdown. Cell viability was analyzed using the CellTiter-Glo luminescent cell viability assay (G7570, Promega, WI, USA) according to the manufacturer’s protocol.

### Protease inhibitors and compound libraries

Nafamostat mesylate (N0959, Tokyo Chemical Industry, Tokyo, Japan), pepstatin A (4397, Peptide institute, Osaka, Japan), bestatin (027-14101, FUJIFILM Wako Pure Chemical), leupeptin (4041, Peptide institute), E-64d (4321-v, Peptide institute), furin inhibitor II (344931, Merck), ilomastat (HY-15768, MedChemExpress, NJ, USA), CTS-1027 (HY-10398, MedChemExpress), marimastat (HY-12169, MedChemExpress), prinomastat hydrochloride (PZ0198, Sigma-Aldrich, MO, USA), UK370106 (2900, Tocris, MN, USA), GW280264X (31388, Cayman, MI, USA), GI254023X (SML0789, Sigma-Aldrich), TAPI-2 (14695, Cayman), MLN-4760 (530616, Merck), BK-1361 (PC-60981, ProbeChem, Shanghai, China), MMP408 (444291, Millipore, MA, USA), MMP2/9 inhibitor I (ab145190, Abcam, Cambridge, UK), and MMP9 inhibitor I (444278, Millipore) were dissolved in dimethyl sulfoxide (DMSO) at a concentration of 10 mM. Validated Compound Library (1,630 clinically approved compounds and 1,885 pharmacologically active compounds) obtained from the Drug Discovery Initiative (The University of Tokyo) was used for compound screening.

### Expression vector construction

To construct expression vectors for ACE2, CD26, and TMPRSS2, genes were cloned into a lentiviral transfer plasmid (CD500B-1, SBI, Palo Alto, CA, USA). Synthetic DNA corresponding to the codon-optimized S gene of SARS-CoV-2 (Wuhan-Hu-1, RefSeq: NC_045512.2), SARS-CoV-2 variants (B.1.1.7, GISAID: EPI_ISL_601443, B.1.351, GenBank: MZ747297.1, B.1.617.1, GISAID: EPI_ISL_1704611, B.1.617.2, GISAID: EPI_ISL_3189054), SARS-CoV (Tor2, RefSeq: NC_004718.3), bat SARS-like coronavirus WIV1 (GenBank: KF367457.1), human coronavirus NL63 (RefSeq: NC_005831.2), and the chimeric S gene (S1, S1/S2 boundary, and S2 domains were derived from either SARS-CoV Tor2 or SARS-Cov-2 Wuhan-Hu-1), and the DNA sequence corresponding to the Flag-tag 5′-GGA GGC <u>GAT TAC AAG GAT GAC GAT GAC AAG</u> TAA-3’ (underline, Flag-tag) at the 3′ end were all generated by Integrated DNA Technologies (IA, USA). Previously described synthetic DNA corresponding to the codon-optimized S gene of a MERS-CoV (EMC 2012, RefSeq: NC_019843.3)[25] with a DNA sequence corresponding to the Flag-tag 5′-GGA GGC GAT TAC AAG GAT GAC GAT GAC AAG TAA-3’ at the 3′ end was used in this study. To construct expression vectors for the S protein, the coding regions were cloned into a lentiviral transfer plasmid (CD500B-1, SBI).

### DSP assay to monitor membrane fusion

DSP1-7 has the structure RL_1–155_-Ser-Gly-Gly-Gly-Gly-GFP_1–156_, while DSP8-11 has the structure Met-GFP_157–231_-Gly-Gly-Gly-Gly-Ser-RL_156–311_. RL and GFP become active only when DSP1-7 associates with DSP-8-11 (S1c Fig). For the DSP assay using 293FT cells, DSP8-11 expressing effector cells expressing S protein and DSP1-7 expressing target cells expressing CD26 or ACE2 alone or together with TMPRSS2 were seeded in 10 cm cell culture plates (4 × 10^6^ cells/10 mL) one day prior to the assay (S1a,b Fig). Cells were treated with 6 μM EnduRen (Promega), a substrate for Renilla luciferase (RL), for 2 h to activate EnduRen. For compound library screening, 0.25 μL of each compound dissolved in DMSO were added to the 384-well plates (Greiner Bioscience, Frickenhausen, Germany). To test the effects of the selected inhibitors, 1 μL of each inhibitor dissolved in DMSO was added to the 384-well plates (Greiner Bioscience). Next, 50 μL of each single cell suspension (effector and target cells) was added to the 384-well plates using a Multidrop dispenser (Thermo Fisher Scientific, MA, USA). After incubation at 37 °C in 5% CO_2_ for 4 h, the RL activity was measured using a Centro xS960 luminometer (Berthold, Bad Wildbad, Germany).

### Western blotting

Western blot analysis was performed as described previously[70]. The primary antibodies used were rabbit anti-ACE2 (1:1000, ab15348, Abcam), rabbit anti-TACE (1:1000, 3976S, Cell Signaling Technology, MA, USA), rabbit anti-ADAM10 (1:1000, 14194S, Cell Signaling Technology), rabbit anti-Flag-tag (1:1000, PM020, MBL, MA, USA), mouse anti-tubulin (1:1000, CP06, Millipore), and mouse anti-VSVM (1:1000, 23H12, Absolute antibody). The Secondly antibodies used were HRP-linked donkey anti-rabbit IgG antibody (NA934; GE Healthcare, Piscataway, NJ, USA) and HRP-linked donkey anti-mouse IgG antibody (NA931V; GE Healthcare). Cell supernatants containing the pseudotype viral particles were centrifuged at 109,000 g or 35 min at 4 °C using a TLA100.3 rotor with an Optima TLX ultracentrifuge (Beckman Coulter, CA, USA), and the pellet was then lysed for western blotting analysis.

### Preparation of pseudotype VSV viral particles and infection experiments

To prepare pseudotype VSV viral particles 293T cells were transfected with an expression plasmid for S, VSV G, or a control expression plasmid using calcium phosphate precipitation. At 16 h post-transfection, the cells were inoculated with a replication-deficient VSV, Δ VSV-Luci, which lacks the VSV G gene and encodes firefly luciferase, at a multiplicity of infection (MOI) of 1, as was described previously[71]. After incubation at 37 °C in 5% CO_2_ for 2 h, the cells were washed with DMEM and further incubated at 37 °C in 5% CO_2_ for 16 h before the supernatants containing the pseudotype viral particles were harvested. Cellular debris was removed from the supernatants using a syringe filter with a 0.45 μm size pore (Millipore). For the infection assay, target cells were seeded in 96-well plates (2 × 10^4^ cells/well) and incubated overnight at 37 °C with 5% CO_2_. The cells were pre-treated with inhibitors for 1 h before infection. Pseudotype viral particles were added to the cells in the presence of the inhibitors. Luciferase activity was measured 16 h post-infection using the Bright-Glo Luciferase Assay System or ONE-Glo Luciferase Assay System (Promega) and Centro xS960 luminometer (Berthold).

### Transfection

siRNA transfection was performed using Lipofectamine RNAiMAX (Thermo Fisher Scientific) according to the manufacturer’s protocol. Cells were seeded in 6-well plates (1.6 × 10^6^ cells/well) with siRNA and Lipofectamine RNAiMAX. Then 24 h after transfection, the cells were seeded in 96-well plates (2 × 10^4^ cells/well), and 48 h after transfection the cells were used for the infection experiments.

### Quantification of intracellular SARS-CoV-2 RNA

Cells were seeded at 5 × 10^4^ cells per well in a 96-well cell culture plate. After an overnight incubation at 37 °C in 5% CO_2_, cells were treated with protease inhibitors for 1 h and added with SARS-CoV-2 at a multiplicity of infection (MOI) of 0.01 for HEC50B and HEC50B-TMPRSS2 cells, and MOI of 0.1 for VeroE6, Calu-3, and A704 cells. After 24 h of incubation at 37 °C in 5% CO_2_, the cells were washed three times with PBS. Cell lysis and cDNA synthesis were performed using SuperPrep II Cell Lysis & RT Kit for qPCR (SCQ-401, TOYOBO, Osaka, Japan) following the manufacturer’s instructions. SARS-CoV-2 RNA was detected using a primer set targeting the SARS-CoV-2 N gene. Quantitative real-time RT-PCR was performed using THUNDERBIRD SYBR qPCR Mix (TOYOBO) at 95 °C for 3 min, followed by 50 cycles of 95 °C for 10 s and 60 °C for 1 min. Fluorescence was detected during the thermal cycling process, and quantification studies were performed using the CFX ConnectTM Real-Time PCR detection system (Bio-Rad, CA, USA). The level of ribosomal protein L13a (*Rpl13a*) mRNA expression in each sample was used to standardize the data. The primer sequences used were 5′-AAA TTT TGG GGA CCA GGA AC-3’ (forward primer) and 5′-TGG CAG CTG TGT AGG TCA AC-3’ (reverse primer) for the SARS-CoV-2 N gene; 5′-TGT TTG ACG GCA TCC CAC-3’ (forward primer) and 5′-CTG TCA CTG CCT GGT ACT TC-3’ (reverse primer) for human *Rpl13a* gene; and 5′-CTC AAG GTT GTG CGT CTG AA-3’ (forward primer) and 5′-CTG TCA CTG CCT GGT ACT TCC A-3’ (reverse primer) for the African green monkey *Rpl13a* gene.

### Immunofluorescence Staining

HEC50B and HEC50B-TMPRSS2 cells were seeded at 1.5 × 10^5^ cells per well in a 24-well cell culture plate. After an overnight incubation at 37 °C in 5% CO_2_, cells were treated with protease inhibitors for 1 h and then with SARS-CoV-2 at an MOI of 1. After 24 h of incubation at 37 °C in 5% CO_2_, cells were fixed with 4% paraformaldehyde in PBS(-) for 10 min at RT and then permeabilized with 0.1% Triton X-100 in PBS(-) for 10 min at RT. Cells were then incubated with anti-SARS-CoV-2 nucleocapsid (1:1000, GTX135357, GeneTex, CA, USA) primary antibody for 16 h at 4 °C and detected with anti-rabbit-Alexa488 (1:200, A11008, Invitrogen, CA, USA) secondary antibodies for 40 min at RT. Cell nuclei were stained with 1 μg/mL Hoechst 33342 (#080-09981, FUJIFILM Wako Pure Chemical). Fluorescent signals were detected using a BZ-X810 fluorescent microscope (Keyence, Osaka, Japan).

### Cytopathicity assay

HEC50B and HEC50B-TMPRSS2 cells were seeded at 1.5 × 10^5^ cells per well in a 24-well cell culture plate. After an overnight incubation at 37 °C in 5% CO_2_, cells were treated with protease inhibitors for 1 h and then with SARS-CoV-2 at an MOI of 1. To maintain the drug concentration, half of the culture supernatant was replaced daily with fresh medium that contained drugs. After incubation at 37 °C in 5% CO_2_ for 3 d, the cells were fixed with 4% paraformaldehyde in PBS for 10 min at 25 °C and stained with 0.2% crystal violet solution for 5 min. After washing four times with water, the wells were air-dried at 25 °C. Ethanol was added to each well to dissolve crystal violet. The absorbance was measured at 595 nm using an iMarkTM Microplate Reader (Bio-Rad).

### Statistical analysis

Statistical analyses were performed in Microsoft Excel 2016 (Microsoft, Redmond, WA, USA) and GraphPad Prism 8 (GraphPad Software, San Diego, CA). Statistically significant differences between the mean values were determined using a two-tailed Student’s t-test. Dunnett’s test and Tukey’s test were used for multiple comparisons. All data represent three independent experiments, and values represent the mean ± standard deviation (s.d.), with a p < 0.05, considered statistically significant.

## Acknowledgements

We thank Yoshihiro Kawaoka for providing the SARS-CoV-2 isolate (UT-NCGM/Human/2020/Tokyo), Robert Whittier for critical reading of our manuscript and Kinuyo Miyazaki for her secretarial assistance.

## Author contributions

M.Y., J.G., and J.I. designed the study; M.Y., J.G., A.K., K.T., and Y.H. performed the experiments; M.Y., J.G., N.K., M.S., K.S., T.A., Y.K., and J.I. analyzed and interpreted the data; and M.Y. and J.I. wrote the manuscript.

## Additional information Financial Disclosure Statement

This work was supported, in part, by the Platform Project for Supporting Drug Discovery and Life Science Research from the Japan Agency for Medical Research and Development (AMED) under Grant Number JP20am0101086 (support number 2834), and by grants-in-aid from the Ministry of Education, Culture, Sports, Science, and Technology, Japan (MEXT; 16H06575 to JI), from the Japanese Society for the Promotion of Science (JSPS; 15K21438 and 18K15235 to MY), from AMED [Program of Japan Initiative for Global Research Network on Infectious Diseases (JGRID) JP20wm0125002 to YK], and from the University of Tokyo (Promoting practical use of measures against coronavirus disease 2019 [COVID-19] to JI). The funders had no role in study design, data collection and analysis, decision to publish, or preparation of the manuscript.

## Supporting information

**S1 Fig. Cell-based membrane-fusion assay for coronavirus S proteins using the DSP reporter.**

**(a**) A method to monitor cell-cell membrane fusion mediated by the S protein of coronaviruses[24, 25]. Effector cells (293FT cells expressing DSP8-11 and S protein) and target cells (293FT cells expressing DSP1-7 and receptor protein with TMPRSS2 for “cell fusion with TMPRSS2” (top) or receptor protein without TMPRSS2 for “cell fusion without TMPRSS2” (bottom)) were co-cultured for 4 h. Both GFP (fluorescence) and RL (luminescence) signals were generated following DSP1-7 and DSP8-11 reassociation upon mixing of the cells during the assay. (**b**) Expression of S proteins in effector cells were detected using an anti-Flag-tag antibody that binds to a Flag-tag on the C-terminus of S proteins (top). Tubulin was used as a control (bottom panel). S0: uncleaved S protein; S2: cleaved S2 domain of the S protein. (**c**) Schematic diagram of split chimeric reporter proteins. DSP1-7 has the structure RL1–155-Ser-Gly-Gly-Gly-Gly-GFP1–156. DSP8-11 has the Met-GFP157–231 -Gly-Gly-Gly-Gly-Ser-RL156–311. As GFP1–156 contains the first seven β sheets, and GFP157–231 contains the remaining four β sheets, the split proteins were called DSP1-7 and DSP8-11, respectively. DSP1-7 and DSP8-11 reassociate efficiently, resulting in the reconstitution of functional RL and GFP to generate luminescent and fluorescent signals, respectively.

**S2 Fig. Control experiments for the DSP assay.**

**(a**) A method to check whether compounds directly inhibit DSP activity without affecting cell-cell fusion[25]. 293FT cells expressing DSP1-7 and DSP8-11 were treated with compounds for 4 h. Measuring RL activities of the preformed DSP1-7/DSP8-11 complex to check whether the compounds directly inhibit RL activities without affecting cell-cell fusion. (**b**) Effect of metalloproteinase inhibitors on RL activity. Relative DSP activity was calculated by normalizing the RL activity for each condition to that of the control assay (DMSO alone; set to 100%). Values are means ± SD (*n* = 3/group).

**S3 Fig. Effects of candidate compounds on the cell-cell fusion and RL activities.**

Effector cells expressing SARS-CoV-2 S were co-cultured with target cells expressing ACE2 alone for the TMPRSS2-independent cell-cell fusion assay (blue) or cells expressing ACE2 with TMPRSS2 for the TMPRSS2-dependent cell-cell fusion assay (red) in the presence of candidate compounds for 4 h. Cells expressing DSP1-7 and DSP8-11 in the presence of candidate compounds for 4 h to determine whether compounds directly inhibit RL activities (purple). Relative DSP activity was calculated by normalizing the RL activity for each condition to that of the control assay (DMSO alone; set to 100%). Values are means ± SD (*n* = 3/group).

**S4 Fig. The metalloproteinase-dependent entry pathway strictly requires both the furin-cleavage site and S2 region of S protein of SARS-CoV-2.**

Effects of the drugs on the entry of S protein-bearing vesicular stomatitis virus (VSV) pseudotype virus. The relative pseudovirus entry was calculated by normalizing the FL activity for each condition to the FL activity of the cells infected with pseudovirus in the presence of DMSO alone, which was set to 100%. Values are means ± SD (*n* = 3/group). ** p < 0.01. Cont: cells infected with pseudovirus without S protein. E-64d: 25 μM E-64d, marima: 1 μM marimastat, nafamo: 10 μM nafamostat. (**a, b**) Effects of E-64d and marimastat on the entry of pseudoviruses bearing SARS-CoV S, SARS-CoV-2 S, MERS-CoV S, or VSV G in A704 (a) and VeroE6 (b) cells. (**c, d**) Effects of E-64d and marimastat on the entry of pseudoviruses bearing chimeric S proteins in VeroE6 cells. (**e**) Effects of E-64d and nafamostat on the entry of pseudoviruses bearing SARS-CoV S, SARS-CoV-2 S, MERS-CoV S, or VSV G in VeroE6-TMPRSS2 cells. (**f, g**) Effects of E-64d and nafamostat on the entry of pseudoviruses bearing chimeric S proteins in VeroE6-TMPRSS2 cells. To establish VeroE6 cells expressing TMPRSS2 (VeroE6-TMPRSS2), recombinant pseudotype lentivirus expressing TMPRSS2 was produced using 293T cells with a VSV-G-expressing plasmid. Cells infected with pseudotype viruses were selected with 300 μg/mL hygromycin for at least 1 week.

**S5 Fig. Patterns of entry pathways were conserved in various variants of SARS-CoV-2.**

**(a**) Expression of WT or mutant SARS-CoV-2 S proteins with mutations precent in B.1.1.7, B.1.351, B.1.617.1 and B.1.617.2 variants in the pseudoviruses. S proteins were detected using an anti-Flag-tag antibody that binds to a Flag-tag on the C-terminus of S proteins (top). Detection of vesicular stomatitis virus matrix protein (VSV M) served as a control (bottom). S0: uncleaved S protein; S2: cleaved S2 domain of the S protein. (**b**) Effects of E-64d and marimastat on the entry of pseudoviruses bearing SARS-CoV-2 S in VeroE6 cells. E-64d: 25 μM E-64d, marima: 1 μM marimastat. (**c**) Effects of nafamostat on the entry of pseudovirus bearing SARS-CoV-2 S in VeroE6-TMPRSS2 cells. (**d**) Effects of E-64d and marimastat on the entry of pseudoviruses bearing SARS-CoV-2 S in HEC50B cells. E-64d: 25 μM E-64d, marima: 1 μM marimastat. (**e**) Effects of marimastat on the entry of pseudoviruses bearing SARS-CoV-2 S in A704 cells. (**f**) Effects of nafamostat on the entry of pseudoviruses bearing SARS-CoV-2 S in Calu-3 cells. The relative pseudovirus entry was calculated by normalizing the FL activity for each condition to the FL activity of cells infected with pseudovirus in the presence of DMSO alone, which was set to 100%. Values are means ± SD (*n* = 3/group in b-f). Data were compared with those obtained from cells infected pseudoviruses bearing variant SARS-CoV-2 S in the presence of DMSO alone. * p < 0.05, ** p < 0.01. Cont: cells infected with a pseudovirus without S protein in (**b-f)**.

**S6 Fig. Expression of WIV1-CoV and HCoV-NL63 S protein in pseudoviruses.**

**(a**) Schematic illustration of C-terminally FLAG-tagged S proteins of WIV1-CoV and HCoV-NL63 and amino acid sequences of the residues around the S1/S2 boundary of the coronaviruses (bottom). Numbers refer to amino acid residues. F: Flag tag. Arginine residues in the S1/S2 cleavage site and furin cleavage motif are highlighted in red. (**b**) Expression of S protein in pseudoviruses S proteins were detected using an anti-Flag-tag antibody that binds to a Flag-tag on the C-terminus of S proteins (top). The detection of VSV M served as a control (bottom). S0: uncleaved S protein; S2: cleaved S2 domain of the S protein.

**S7 Fig. Effects of drugs on cell viabilities.**

**(a-c)** VeroE6 (a), HEC50B (b), and A704 (c) cells were treated with various drugs, and cell viability was analyzed using Celltiter-Glo 24 h after the treatment. The relative cell viability was calculated by normalizing the FL activity for each condition to the FL activity of the cells in the presence of DMSO alone, which was set to 100%. Values are means ± SD (*n* = 3/group). (**d, e**) HEC50B (d), and HEC50B-TMPRSS2 (e) cells were treated with various drugs for 3 days. Half of the culture supernatant was replaced daily with fresh medium containing the drugs. The relative cell viability was calculated by normalizing the FL activity for each condition to the FL activity of cells in the presence of DMSO alone, which was set to 100%. Values are means ± SD (*n* = 6/group).

**S8 Fig. Patterns of the entry pathways of the pseudovirus bearing SARS-CoV-2 S were not affected by the ADAM17 knockout in the HEC50B cells.**

**(a**) Effect of the ADAM17 knockout on ACE2 (top), ADAM17 (middle), and tubulin (bottom). (**b**) Effect of the ADAM17 knockout on the entry of the pseudoviruses bearing SARS-CoV-2 S. Values are means ± SD (*n* = 3/group). ** p < 0.01. (**c**) Effect of the ADAM17 knockout on the patterns of the entry pathways of SARS-CoV-2 S pseudovirus in HEC50B cells. The relative pseudovirus entry was calculated by normalizing the FL activity for each condition to the FL activity of cells infected with pseudovirus in the presence of DMSO alone, which was set to 100%. Values are means ± SD (*n* = 3/group). * p < 0.05, ** p < 0.01. E-64d: 25 μM E-64d, marima: 1 μM marimastat. To establish the ADAM17-knockout HEC50B cells, lentiviruses were produced by transfecting the lentiCRISPRv2 vector (#52961 Addgene, MA, USA) with the following gRNA sequences. The gRNA sequences used were 5′-GCG AGG TAT TCG GCT CCG CG-3’ (Cont #1), 5′-GCT TTC ACG GAG GTT CGA CG-3’ (Cont #2) and 5′-ATG TTG CAG TTC GGC TCG AT-3’ (Cont #3) for the control experiments, and 5′-AAC GTT CAG TAC TTG ATG TC-3’ (ADAM10 #1) and 5′-GGA CTT CTT CAC TGG ACA CG-3’ (ADAM10 #2) and 5′-CTT AAG GTG AGC CTG ACT CT-3’ (ADAM10 #3) for the establishment of ADAM17-knockout cells. Pooled HEC50B cells infected with pseudotype viruses were selected with 1 μg/mL puromycin for 1 week.

**S9 Fig. Effects of drugs on SARS-CoV-2 infection.**

**(a**) Effects of the nafamostat on the SARS-CoV-2 infection in Calu-3, HEC50B, A704, and VeroE6 cells. Values are means ± SD (*n* = 3/group). ** p < 0.01. (**b**) Effects of the E-64d on the SARS-CoV-2 infection in HEC50B, A704, and VeroE6 cells. Values are means ± SD (*n* = 3/group). * p < 0.05, ** p < 0.01. (**c**) Effects of the NH_4_Cl on the SARS-CoV-2 infection in HEC50B cells. Values are means (*n* = 2/group). The relative amount of viral RNA in the cells was normalized to cellular *Rpl13a* mRNA expression in (a-c).

**S10 Fig. The metalloproteinase-dependent entry pathway of authentic SARS-CoV-2 is involved in syncytia formation.**

Phase contrast images of syncytia formation 24 h after SARS-CoV-2 infection in the presence of inhibitors. Red arrowheads indicate syncytia formation Scale bars, 100 μm.

## References

1. Zhou P, Yang XL, Wang XG, Hu B, Zhang L, Zhang W, et al. A pneumonia outbreak associated with a new coronavirus of probable bat origin. Nature. 2020;579(7798):270–3. Epub 2020/02/03. doi: 10.1038/s41586-020-2012-7. PubMed PMID: 32015507; PubMed Central PMCID: PMCPMC7095418.

2. Zhong NS, Zheng BJ, Li YM, Poon, Xie ZH, Chan KH, et al. Epidemiology and cause of severe acute respiratory syndrome (SARS) in Guangdong, People’s Republic of China, in February, 2003. Lancet. 2003;362(9393):1353–8. doi: 10.1016/s0140-6736(03)14630-2. PubMed PMID: 14585636; PubMed Central PMCID: PMCPMC7112415.

3. Drosten C, Günther S, Preiser W, van der Werf S, Brodt HR, Becker S, et al. Identification of a novel coronavirus in patients with severe acute respiratory syndrome. N Engl J Med. 2003;348(20):1967–76. Epub 20030410. doi: 10.1056/NEJMoa030747. PubMed PMID: 12690091.

4. Zaki AM, van Boheemen S, Bestebroer TM, Osterhaus AD, Fouchier RA. Isolation of a novel coronavirus from a man with pneumonia in Saudi Arabia. N Engl J Med. 2012;367(19):1814–20. doi: 10.1056/NEJMoa1211721. PubMed PMID: 23075143.

5. Harrison AG, Lin T, Wang P. Mechanisms of SARS-CoV-2 Transmission and Pathogenesis. Trends Immunol. 2020;41(12):1100–15. Epub 20201014. doi: 10.1016/j.it.2020.10.004. PubMed PMID: 33132005; PubMed Central PMCID: PMCPMC7556779.

6. Hu B, Guo H, Zhou P, Shi ZL. Characteristics of SARS-CoV-2 and COVID-19. Nat Rev Microbiol. 2021;19(3):141–54. Epub 20201006. doi: 10.1038/s41579-020-00459-7. PubMed PMID: 33024307; PubMed Central PMCID: PMCPMC7537588.

7. Tregoning JS, Flight KE, Higham SL, Wang Z, Pierce BF. Progress of the COVID-19 vaccine effort: viruses, vaccines and variants versus efficacy, effectiveness and escape. Nat Rev Immunol. 2021;21(10):626–36. Epub 20210809. doi: 10.1038/s41577-021-00592-1. PubMed PMID: 34373623; PubMed Central PMCID: PMCPMC8351583.

8. Pritchard E, Matthews PC, Stoesser N, Eyre DW, Gethings O, Vihta KD, et al. Impact of vaccination on new SARS-CoV-2 infections in the United Kingdom. Nat Med. 2021;27(8):1370–8. Epub 20210609. doi: 10.1038/s41591-021-01410-w. PubMed PMID: 34108716; PubMed Central PMCID: PMCPMC8363500.

9. Planas D, Veyer D, Baidaliuk A, Staropoli I, Guivel-Benhassine F, Rajah MM, et al. Reduced sensitivity of SARS-CoV-2 variant Delta to antibody neutralization. Nature. 2021;596(7871):276–80. Epub 20210708. doi: 10.1038/s41586-021-03777-9. PubMed PMID: 34237773.

10. Tao K, Tzou PL, Nouhin J, Gupta RK, de Oliveira T, Kosakovsky Pond SL, et al. The biological and clinical significance of emerging SARS-CoV-2 variants. Nat Rev Genet. 2021. Epub 20210917. doi: 10.1038/s41576-021-00408-x. PubMed PMID: 34535792; PubMed Central PMCID: PMCPMC8447121.

11. Uriu K, Kimura I, Shirakawa K, Takaori-Kondo A, Nakada TA, Kaneda A, et al. Neutralization of the SARS-CoV-2 Mu Variant by Convalescent and Vaccine Serum. N Engl J Med. 2021. Epub 20211103. doi: 10.1056/NEJMc2114706. PubMed PMID: 34731554.

12. Mei M, Tan X. Current Strategies of Antiviral Drug Discovery for COVID-19. Front Mol Biosci. 2021;8:671263. Epub 20210513. doi: 10.3389/fmolb.2021.671263. PubMed PMID: 34055887; PubMed Central PMCID: PMCPMC8155633.

13. Rando HM, Wellhausen N, Ghosh S, Lee AJ, Dattoli AA, Hu F, et al. Identification and Development of Therapeutics for COVID-19. mSystems. 2021:e0023321. Epub 20211102. doi: 10.1128/mSystems.00233-21. PubMed PMID: 34726496; PubMed Central PMCID: PMCPMC8562484.

14. Jackson CB, Farzan M, Chen B, Choe H. Mechanisms of SARS-CoV-2 entry into cells. Nat Rev Mol Cell Biol. 2021. Epub 20211005. doi: 10.1038/s41580-021-00418-x. PubMed PMID: 34611326; PubMed Central PMCID: PMCPMC8491763.

15. Hoffmann M, Kleine-Weber H, Pöhlmann S. A Multibasic Cleavage Site in the Spike Protein of SARS-CoV-2 Is Essential for Infection of Human Lung Cells. Mol Cell. 2020;78(4):779–84.e5. Epub 20200501. doi: 10.1016/j.molcel.2020.04.022. PubMed PMID: 32362314; PubMed Central PMCID: PMCPMC7194065.

16. Shang J, Wan Y, Luo C, Ye G, Geng Q, Auerbach A, et al. Cell entry mechanisms of SARS-CoV-2. Proc Natl Acad Sci U S A. 2020;117(21):11727–34. Epub 2020/05/06. doi: 10.1073/pnas.2003138117. PubMed PMID: 32376634.

17. Yan R, Zhang Y, Li Y, Xia L, Guo Y, Zhou Q. Structural basis for the recognition of SARS-CoV-2 by full-length human ACE2. Science. 2020;367(6485):1444–8. Epub 20200304. doi: 10.1126/science.abb2762. PubMed PMID: 32132184; PubMed Central PMCID: PMCPMC7164635.

18. Lan J, Ge J, Yu J, Shan S, Zhou H, Fan S, et al. Structure of the SARS-CoV-2 spike receptor-binding domain bound to the ACE2 receptor. Nature. 2020;581(7807):215–20. Epub 20200330. doi: 10.1038/s41586-020-2180-5. PubMed PMID: 32225176.

19. Hoffmann M, Kleine-Weber H, Schroeder S, Krüger N, Herrler T, Erichsen S, et al. SARS-CoV-2 Cell Entry Depends on ACE2 and TMPRSS2 and Is Blocked by a Clinically Proven Protease Inhibitor. Cell. 2020;181(2):271–80.e8. Epub 2020/03/05. doi: 10.1016/j.cell.2020.02.052. PubMed PMID: 32142651; PubMed Central PMCID: PMCPMC7102627.

20. Zhao MM, Yang WL, Yang FY, Zhang L, Huang WJ, Hou W, et al. Cathepsin L plays a key role in SARS-CoV-2 infection in humans and humanized mice and is a promising target for new drug development. Signal Transduct Target Ther. 2021;6(1):134. Epub 20210327. doi: 10.1038/s41392-021-00558-8. PubMed PMID: 33774649; PubMed Central PMCID: PMCPMC7997800.

21. Murgolo N, Therien AG, Howell B, Klein D, Koeplinger K, Lieberman LA, et al. SARS-CoV-2 tropism, entry, replication, and propagation: Considerations for drug discovery and development. PLoS Pathog. 2021;17(2):e1009225. Epub 2021/02/17. doi: 10.1371/journal.ppat.1009225. PubMed PMID: 33596266; PubMed Central PMCID: PMCPMC7888651.

22. Koch J, Uckeley ZM, Doldan P, Stanifer M, Boulant S, Lozach PY. TMPRSS2 expression dictates the entry route used by SARS-CoV-2 to infect host cells. EMBO J. 2021;40(16):e107821. Epub 20210713. doi: 10.15252/embj.2021107821. PubMed PMID: 34159616; PubMed Central PMCID: PMCPMC8365257.

23. Bestle D, Heindl MR, Limburg H, Van Lam van T, Pilgram O, Moulton H, et al. TMPRSS2 and furin are both essential for proteolytic activation of SARS-CoV-2 in human airway cells. Life Sci Alliance. 2020;3(9). Epub 20200723. doi: 10.26508/lsa.202000786. PubMed PMID: 32703818; PubMed Central PMCID: PMCPMC7383062.

24. Yamamoto M, Kiso M, Sakai-Tagawa Y, Iwatsuki-Horimoto K, Imai M, Takeda M, et al. The Anticoagulant Nafamostat Potently Inhibits SARS-CoV-2 S Protein-Mediated Fusion in a Cell Fusion Assay System and Viral Infection In Vitro in a Cell-Type-Dependent Manner. Viruses. 2020;12(6). Epub 2020/06/10. doi: 10.3390/v12060629. PubMed PMID: 32532094.

25. Yamamoto M, Matsuyama S, Li X, Takeda M, Kawaguchi Y, Inoue JI, et al. Identification of Nafamostat as a Potent Inhibitor of Middle East Respiratory Syndrome Coronavirus S Protein-Mediated Membrane Fusion Using the Split-Protein-Based Cell-Cell Fusion Assay. Antimicrob Agents Chemother. 2016;60(11):6532–9. Epub 2016/08/24. doi: 10.1128/aac.01043-16. PubMed PMID: 27550352; PubMed Central PMCID: PMCPMC5075056.

26. Ishikawa H, Meng F, Kondo N, Iwamoto A, Matsuda Z. Generation of a dual-functional split-reporter protein for monitoring membrane fusion using self-associating split GFP. Protein Eng Des Sel. 2012;25(12):813–20. doi: 10.1093/protein/gzs051. PubMed PMID: 22942393.

27. Wojtowicz-Praga S, Torri J, Johnson M, Steen V, Marshall J, Ness E, et al. Phase I trial of Marimastat, a novel matrix metalloproteinase inhibitor, administered orally to patients with advanced lung cancer. J Clin Oncol. 1998;16(6):2150–6. doi: 10.1200/JCO.1998.16.6.2150. PubMed PMID: 9626215.

28. Hande KR, Collier M, Paradiso L, Stuart-Smith J, Dixon M, Clendeninn N, et al. Phase I and pharmacokinetic study of prinomastat, a matrix metalloprotease inhibitor. Clin Cancer Res. 2004;10(3):909–15. doi: 10.1158/1078-0432.ccr-0981-3. PubMed PMID: 14871966.

29. Standard of Care (SOC) With or Without CTS-1027 in Hepatitis C (HCV) Null-Responders https://clinicaltrials.gov/ct2/show/results/NCT01273064.

30. Vandenbroucke RE, Dejonckheere E, Libert C. A therapeutic role for matrix metalloproteinase inhibitors in lung diseases? Eur Respir J. 2011;38(5):1200–14. Epub 20110609. doi: 10.1183/09031936.00027411. PubMed PMID: 21659416.

31. Jacobsen JA, Major Jourden JL, Miller MT, Cohen SM. To bind zinc or not to bind zinc: an examination of innovative approaches to improved metalloproteinase inhibition. Biochim Biophys Acta. 2010;1803(1):72–94. Epub 20090825. doi: 10.1016/j.bbamcr.2009.08.006. PubMed PMID: 19712708.

32. Madoux F, Dreymuller D, Pettiloud JP, Santos R, Becker-Pauly C, Ludwig A, et al. Discovery of an enzyme and substrate selective inhibitor of ADAM10 using an exosite-binding glycosylated substrate. Sci Rep. 2016;6(1):11. Epub 20161205. doi: 10.1038/s41598-016-0013-4. PubMed PMID: 28442704; PubMed Central PMCID: PMCPMC5431342.

33. Liechti FD, Bächtold F, Grandgirard D, Leppert D, Leib SL. The matrix metalloproteinase inhibitor RS-130830 attenuates brain injury in experimental pneumococcal meningitis. J Neuroinflammation. 2015;12:43. Epub 20150304. doi: 10.1186/s12974-015-0257-0. PubMed PMID: 25890041; PubMed Central PMCID: PMCPMC4352253.

34. Hundhausen C, Misztela D, Berkhout TA, Broadway N, Saftig P, Reiss K, et al. The disintegrin-like metalloproteinase ADAM10 is involved in constitutive cleavage of CX3CL1 (fractalkine) and regulates CX3CL1-mediated cell-cell adhesion. Blood. 2003;102(4):1186–95. Epub 20030424. doi: 10.1182/blood-2002-12-3775. PubMed PMID: 12714508.

35. Ludwig A, Hundhausen C, Lambert MH, Broadway N, Andrews RC, Bickett DM, et al. Metalloproteinase inhibitors for the disintegrin-like metalloproteinases ADAM10 and ADAM17 that differentially block constitutive and phorbol ester-inducible shedding of cell surface molecules. Comb Chem High Throughput Screen. 2005;8(2):161–71. doi: 10.2174/1386207053258488. PubMed PMID: 15777180.

36. Black RA, Rauch CT, Kozlosky CJ, Peschon JJ, Slack JL, Wolfson MF, et al. A metalloproteinase disintegrin that releases tumour-necrosis factor-alpha from cells. Nature. 1997;385(6618):729–33. doi: 10.1038/385729a0. PubMed PMID: 9034190.

37. Schlomann U, Koller G, Conrad C, Ferdous T, Golfi P, Garcia AM, et al. ADAM8 as a drug target in pancreatic cancer. Nat Commun. 2015;6:6175. Epub 20150128. doi: 10.1038/ncomms7175. PubMed PMID: 25629724; PubMed Central PMCID: PMCPMC5014123.

38. Tamura Y, Watanabe F, Nakatani T, Yasui K, Fuji M, Komurasaki T, et al. Highly selective and orally active inhibitors of type IV collagenase (MMP-9 and MMP-2): N-sulfonylamino acid derivatives. J Med Chem. 1998;41(4):640–9. doi: 10.1021/jm9707582. PubMed PMID: 9484512.

39. Fray MJ, Dickinson RP, Huggins JP, Occleston NL. A potent, selective inhibitor of matrix metalloproteinase-3 for the topical treatment of chronic dermal ulcers. J Med Chem. 2003;46(16):3514–25. doi: 10.1021/jm0308038. PubMed PMID: 12877590.

40. Levin JI, Chen J, Du M, Hogan M, Kincaid S, Nelson FC, et al. The discovery of anthranilic acid-based MMP inhibitors. Part 2: SAR of the 5-position and P1(1) groups. Bioorg Med Chem Lett. 2001;11(16):2189–92. doi: 10.1016/s0960-894x(01)00419-x. PubMed PMID: 11514167.

41. Dales NA, Gould AE, Brown JA, Calderwood EF, Guan B, Minor CA, et al. Substrate-based design of the first class of angiotensin-converting enzyme-related carboxypeptidase (ACE2) inhibitors. J Am Chem Soc. 2002;124(40):11852–3. doi: 10.1021/ja0277226. PubMed PMID: 12358520.

42. Yeung ML, Teng JLL, Jia L, Zhang C, Huang C, Cai JP, et al. Soluble ACE2-mediated cell entry of SARS-CoV-2 via interaction with proteins related to the renin-angiotensin system. Cell. 2021;184(8):2212–28.e12. Epub 20210302. doi: 10.1016/j.cell.2021.02.053. PubMed PMID: 33713620; PubMed Central PMCID: PMCPMC7923941.

43. Rodríguez E, Everitt E. Adenovirus uncoating and nuclear establishment are not affected by weak base amines. J Virol. 1996;70(6):3470–7. doi: 10.1128/JVI.70.6.3470-3477.1996. PubMed PMID: 8648679; PubMed Central PMCID: PMCPMC190220.

44. Zhang Z, Zheng Y, Niu Z, Zhang B, Wang C, Yao X, et al. SARS-CoV-2 spike protein dictates syncytium-mediated lymphocyte elimination. Cell Death Differ. 2021;28(9):2765–77. Epub 20210420. doi: 10.1038/s41418-021-00782-3. PubMed PMID: 33879858; PubMed Central PMCID: PMCPMC8056997.

45. Bussani R, Schneider E, Zentilin L, Collesi C, Ali H, Braga L, et al. Persistence of viral RNA, pneumocyte syncytia and thrombosis are hallmarks of advanced COVID-19 pathology. EBioMedicine. 2020;61:103104. Epub 20201103. doi: 10.1016/j.ebiom.2020.103104. PubMed PMID: 33158808; PubMed Central PMCID: PMCPMC7677597.

46. Hoffmann M, Schroeder S, Kleine-Weber H, Müller MA, Drosten C, Pöhlmann S. Nafamostat mesylate blocks activation of SARS-CoV-2: New treatment option for COVID-19. Antimicrob Agents Chemother. 2020. Epub 2020/04/20. doi: 10.1128/AAC.00754-20. PubMed PMID: 32312781.

47. Fan C, Lu W, Li K, Ding Y, Wang J. ACE2 Expression in Kidney and Testis May Cause Kidney and Testis Infection in COVID-19 Patients. Front Med (Lausanne). 2020;7:563893. Epub 20210113. doi: 10.3389/fmed.2020.563893. PubMed PMID: 33521006; PubMed Central PMCID: PMCPMC7838217.

48. Liu J, Li Y, Liu Q, Yao Q, Wang X, Zhang H, et al. SARS-CoV-2 cell tropism and multiorgan infection. Cell Discov. 2021;7(1):17. Epub 20210323. doi: 10.1038/s41421-021-00249-2. PubMed PMID: 33758165; PubMed Central PMCID: PMCPMC7987126.

49. Reis FM, Bouissou DR, Pereira VM, Camargos AF, dos Reis AM, Santos RA. Angiotensin-(1-7), its receptor Mas, and the angiotensin-converting enzyme type 2 are expressed in the human ovary. Fertil Steril. 2011;95(1):176–81. Epub 20100801. doi: 10.1016/j.fertnstert.2010.06.060. PubMed PMID: 20674894.

50. Jing Y, Run-Qian L, Hao-Ran W, Hao-Ran C, Ya-Bin L, Yang G, et al. Potential influence of COVID-19/ACE2 on the female reproductive system. Mol Hum Reprod. 2020;26(6):367–73. doi: 10.1093/molehr/gaaa030. PubMed PMID: 32365180; PubMed Central PMCID: PMCPMC7239105.

51. Henarejos-Castillo I, Sebastian-Leon P, Devesa-Peiro A, Pellicer A, Diaz-Gimeno P. SARS-CoV-2 infection risk assessment in the endometrium: viral infection-related gene expression across the menstrual cycle. Fertil Steril. 2020;114(2):223–32. Epub 20200617. doi: 10.1016/j.fertnstert.2020.06.026. PubMed PMID: 32641214; PubMed Central PMCID: PMCPMC7298504.

52. Puelles VG, Lütgehetmann M, Lindenmeyer MT, Sperhake JP, Wong MN, Allweiss L, et al. Multiorgan and Renal Tropism of SARS-CoV-2. N Engl J Med. 2020;383(6):590–2. Epub 2020/05/13. doi: 10.1056/NEJMc2011400. PubMed PMID: 32402155; PubMed Central PMCID: PMCPMC7240771.

53. Diao B, Wang C, Wang R, Feng Z, Zhang J, Yang H, et al. Human kidney is a target for novel severe acute respiratory syndrome coronavirus 2 infection. Nat Commun. 2021;12(1):2506. Epub 20210504. doi: 10.1038/s41467-021-22781-1. PubMed PMID: 33947851; PubMed Central PMCID: PMCPMC8096808.

54. Johnson BA, Xie X, Bailey AL, Kalveram B, Lokugamage KG, Muruato A, et al. Loss of furin cleavage site attenuates SARS-CoV-2 pathogenesis. Nature. 2021;591(7849):293–9. Epub 20210125. doi: 10.1038/s41586-021-03237-4. PubMed PMID: 33494095; PubMed Central PMCID: PMCPMC8175039.

55. Peacock TP, Goldhill DH, Zhou J, Baillon L, Frise R, Swann OC, et al. The furin cleavage site in the SARS-CoV-2 spike protein is required for transmission in ferrets. Nat Microbiol. 2021;6(7):899–909. Epub 20210427. doi: 10.1038/s41564-021-00908-w. PubMed PMID: 33907312.

56. Benton DJ, Wrobel AG, Xu P, Roustan C, Martin SR, Rosenthal PB, et al. Receptor binding and priming of the spike protein of SARS-CoV-2 for membrane fusion. Nature. 2020;588(7837):327–30. Epub 20200917. doi: 10.1038/s41586-020-2772-0. PubMed PMID: 32942285; PubMed Central PMCID: PMCPMC7116727.

57. Stevens CS, Oguntuyo KY, Lee B. Proteases and variants: context matters for SARS-CoV-2 entry assays. Curr Opin Virol. 2021;50:49–58. Epub 20210724. doi: 10.1016/j.coviro.2021.07.004. PubMed PMID: 34365113; PubMed Central PMCID: PMCPMC8302850.

58. Harvey WT, Carabelli AM, Jackson B, Gupta RK, Thomson EC, Harrison EM, et al. SARS-CoV-2 variants, spike mutations and immune escape. Nat Rev Microbiol. 2021;19(7):409–24. Epub 20210601. doi: 10.1038/s41579-021-00573-0. PubMed PMID: 34075212; PubMed Central PMCID: PMCPMC8167834.

59. 59. The human protein atlas - Tissue expression of ADAM10 https://www.proteinatlas.org/ENSG00000137845-ADAM10/tissue.

60. 60. The human protein atlas - Cell type atlas https://www.proteinatlas.org/ENSG00000137845-ADAM10/celltype.

61. Caescu CI, Jeschke GR, Turk BE. Active-site determinants of substrate recognition by the metalloproteinases TACE and ADAM10. Biochem J. 2009;424(1):79–88. Epub 20091023. doi: 10.1042/BJ20090549. PubMed PMID: 19715556; PubMed Central PMCID: PMCPMC2774824.

62. Carapito R, Li R, Helms J, Carapito C, Gujja S, Rolli V, et al. Identification of driver genes for critical forms of COVID-19 in a deeply phenotyped young patient cohort. Sci Transl Med. 2021:eabj7521. Epub 20211026. doi: 10.1126/scitranslmed.abj7521. PubMed PMID: 34698500.

63. Daniloski Z, Jordan TX, Wessels HH, Hoagland DA, Kasela S, Legut M, et al. Identification of Required Host Factors for SARS-CoV-2 Infection in Human Cells. Cell. 2021;184(1):92–105.e16. Epub 20201024. doi: 10.1016/j.cell.2020.10.030. PubMed PMID: 33147445; PubMed Central PMCID: PMCPMC7584921.

64. Gunst JD, Staerke NB, Pahus MH, Kristensen LH, Bodilsen J, Lohse N, et al. Efficacy of the TMPRSS2 inhibitor camostat mesilate in patients hospitalized with Covid-19-a double-blind randomized controlled trial. EClinicalMedicine. 2021;35:100849. Epub 20210422. doi: 10.1016/j.eclinm.2021.100849. PubMed PMID: 33903855; PubMed Central PMCID: PMCPMC8060682.

65. Boulware DR, Pullen MF, Bangdiwala AS, Pastick KA, Lofgren SM, Okafor EC, et al. A Randomized Trial of Hydroxychloroquine as Postexposure Prophylaxis for Covid-19. N Engl J Med. 2020;383(6):517–25. Epub 20200603. doi: 10.1056/NEJMoa2016638. PubMed PMID: 32492293; PubMed Central PMCID: PMCPMC7289276.

66. Self WH, Semler MW, Leither LM, Casey JD, Angus DC, Brower RG, et al. Effect of Hydroxychloroquine on Clinical Status at 14 Days in Hospitalized Patients With COVID-19: A Randomized Clinical Trial. JAMA. 2020;324(21):2165–76. doi: 10.1001/jama.2020.22240. PubMed PMID: 33165621; PubMed Central PMCID: PMCPMC7653542.

67. Matsuyama S, Nao N, Shirato K, Kawase M, Saito S, Takayama I, et al. Enhanced isolation of SARS-CoV-2 by TMPRSS2-expressing cells. Proc Natl Acad Sci U S A. 2020;117(13):7001–3. Epub 2020/03/12. doi: 10.1073/pnas.2002589117. PubMed PMID: 32165541; PubMed Central PMCID: PMCPMC7132130.

68. Wang H, Li X, Nakane S, Liu S, Ishikawa H, Iwamoto A, et al. Co-expression of foreign proteins tethered to HIV-1 envelope glycoprotein on the cell surface by introducing an intervening second membrane-spanning domain. PLoS One. 2014;9(5):e96790. doi: 10.1371/journal.pone.0096790. PubMed PMID: 24804933; PubMed Central PMCID: PMCPMC4013048.

69. Imai M, Iwatsuki-Horimoto K, Hatta M, Loeber S, Halfmann PJ, Nakajima N, et al. Syrian hamsters as a small animal model for SARS-CoV-2 infection and countermeasure development. Proc Natl Acad Sci U S A. 2020;117(28):16587–95. Epub 20200622. doi: 10.1073/pnas.2009799117. PubMed PMID: 32571934; PubMed Central PMCID: PMCPMC7368255.

70. Yamamoto M, Abe C, Wakinaga S, Sakane K, Yumiketa Y, Taguchi Y, et al. TRAF6 maintains mammary stem cells and promotes pregnancy-induced mammary epithelial cell expansion. Commun Biol. 2019;2:292. Epub 2019/08/06. doi: 10.1038/s42003-019-0547-7. PubMed PMID: 31396572; PubMed Central PMCID: PMCPMC6684589.

71. Tani H, Shiokawa M, Kaname Y, Kambara H, Mori Y, Abe T, et al. Involvement of ceramide in the propagation of Japanese encephalitis virus. J Virol. 2010;84(6):2798–807. Epub 20100106. doi: 10.1128/JVI.02499-09. PubMed PMID: 20053738; PubMed Central PMCID: PMCPMC2826033.

