## Supporting information for "Metalloproteinase-dependent and TMPRSS2-independnt cell surface entry pathway of SARS-CoV-2 requires the furin-cleavage site and the S2 domain of spike protein"

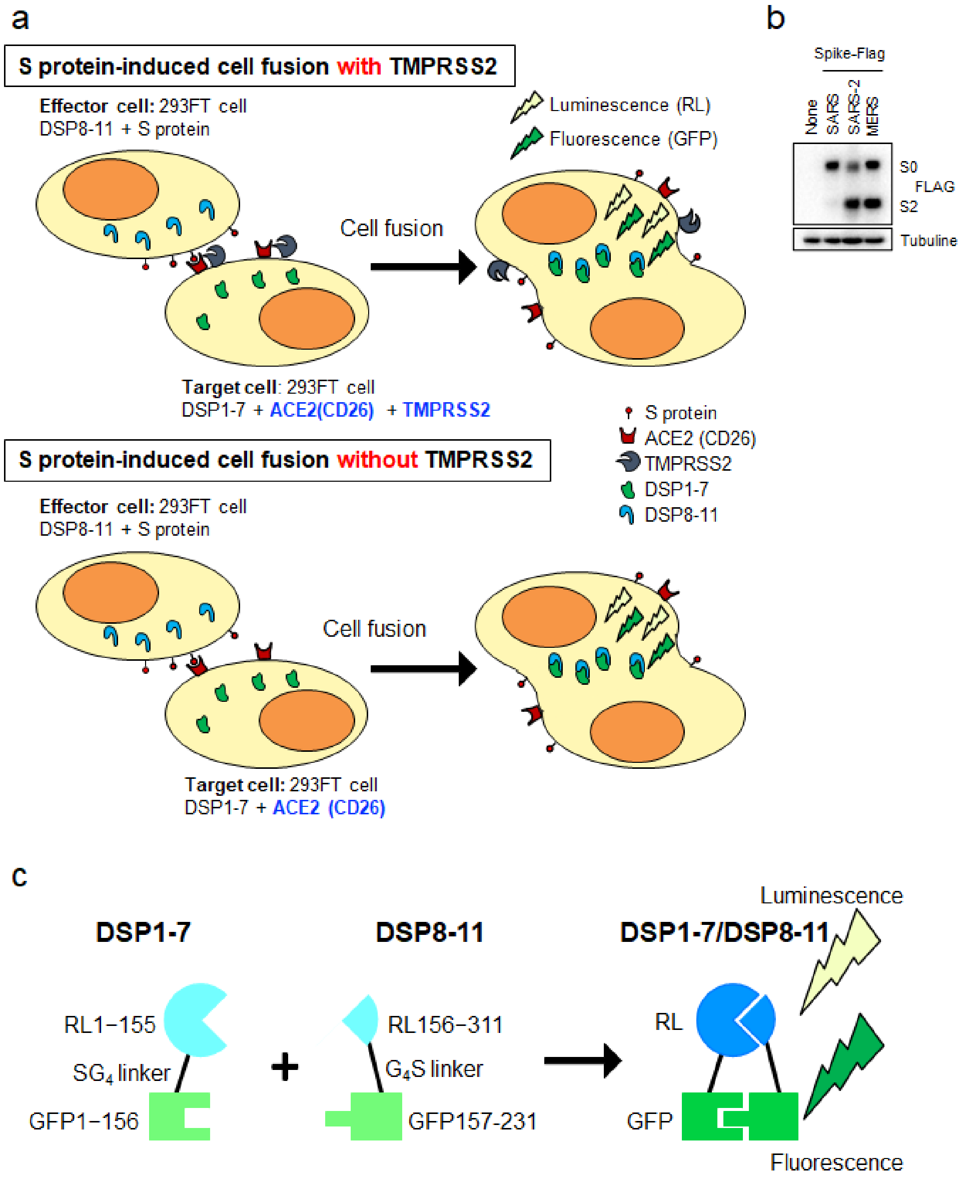


**S1 Fig. Cell-based membrane-fusion assay for coronavirus S proteins using the DSP reporter.**

(**a**) A method to monitor cell-cell membrane fusion mediated by the S protein of coronaviruses[24, 25]. Effector cells (293FT cells expressing DSP8-11 and S protein) and target cells (293FT cells expressing DSP1-7 and receptor protein with TMPRSS2 for "cell fusion with TMPRSS2" (top) or receptor protein without TMPRSS2 for "cell fusion without TMPRSS2" (bottom)) were co-cultured for 4 h. Both GFP (fluorescence) and RL (luminescence) signals were generated following DSP1-7 and DSP8-11 reassociation upon mixing of the cells during the assay. (**b**) Expression of S proteins in effector cells were detected using an anti-Flag-tag antibody that binds to a Flag-tag on the C-terminus of S proteins (top). Tubulin was used as a control (bottom panel). S0: uncleaved S protein; S2: cleaved S2 domain of the S protein. (**c**) Schematic diagram of split chimeric reporter proteins. DSP1-7 has the structure RL1–155-Ser-Gly-Gly-Gly-Gly-GFP1–156. DSP8-11 has the Met- GFP157–231 -Gly-Gly-Gly-Gly-Ser- RL156–311. As GFP1–156 contains the first seven β sheets, and GFP157–231 contains the remaining four β sheets, the split proteins were called DSP1-7 and DSP8-11, respectively. DSP1-7 and DSP8-11 reassociate efficiently, resulting in the reconstitution of functional RL and GFP to generate luminescent and fluorescent signals, respectively.

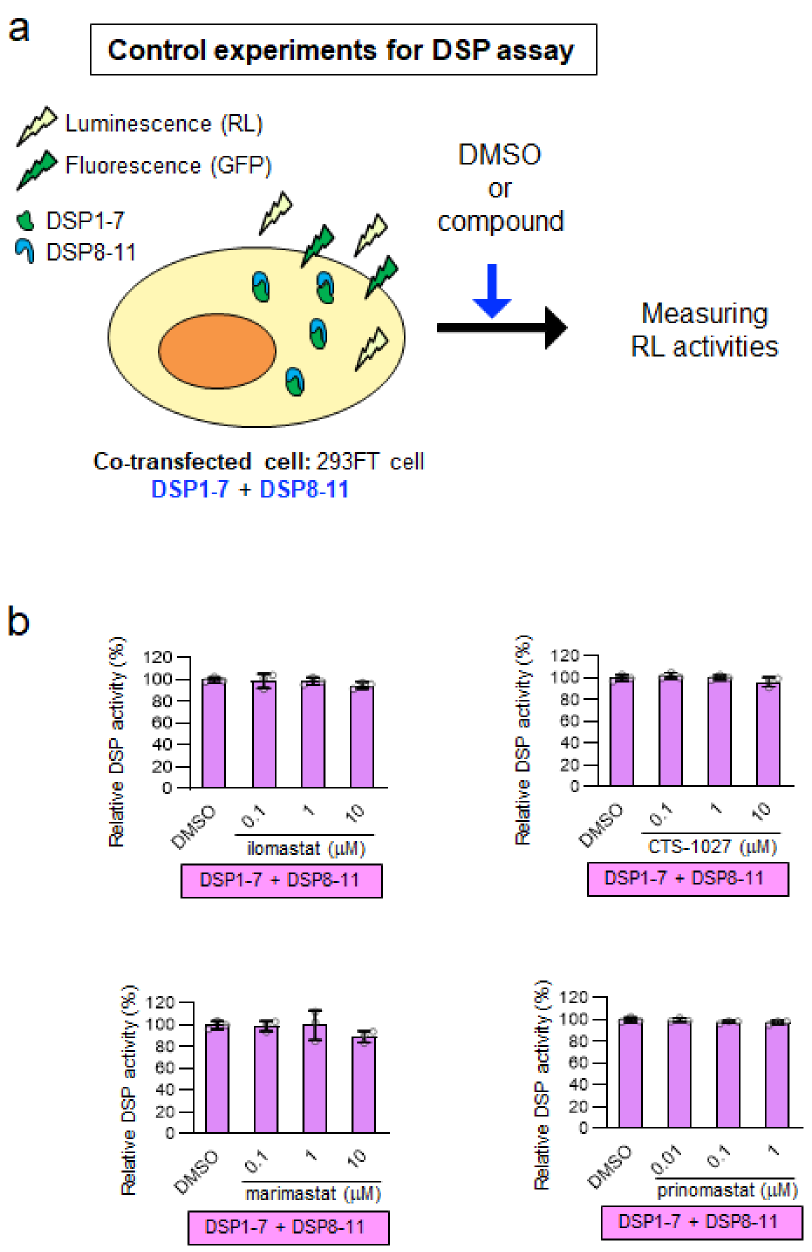


**S2 Fig. Control experiments for the DSP assay.**

(**a**) A method to check whether compounds directly inhibit DSP activity without affecting cell-cell fusion[25]. 293FT cells expressing DSP1-7 and DSP8-11 were treated with compounds for 4 h. Measuring RL activities of the preformed DSP1-7/DSP8-11 complex to check whether the compounds directly inhibit RL activities without affecting cell-cell fusion. (**b**) Effect of metalloproteinase inhibitors on RL activity. Relative DSP activity was calculated by normalizing the RL activity for each condition to that of the control assay (DMSO alone; set to 100%). Values are means ± SD (*n* = 3/group).

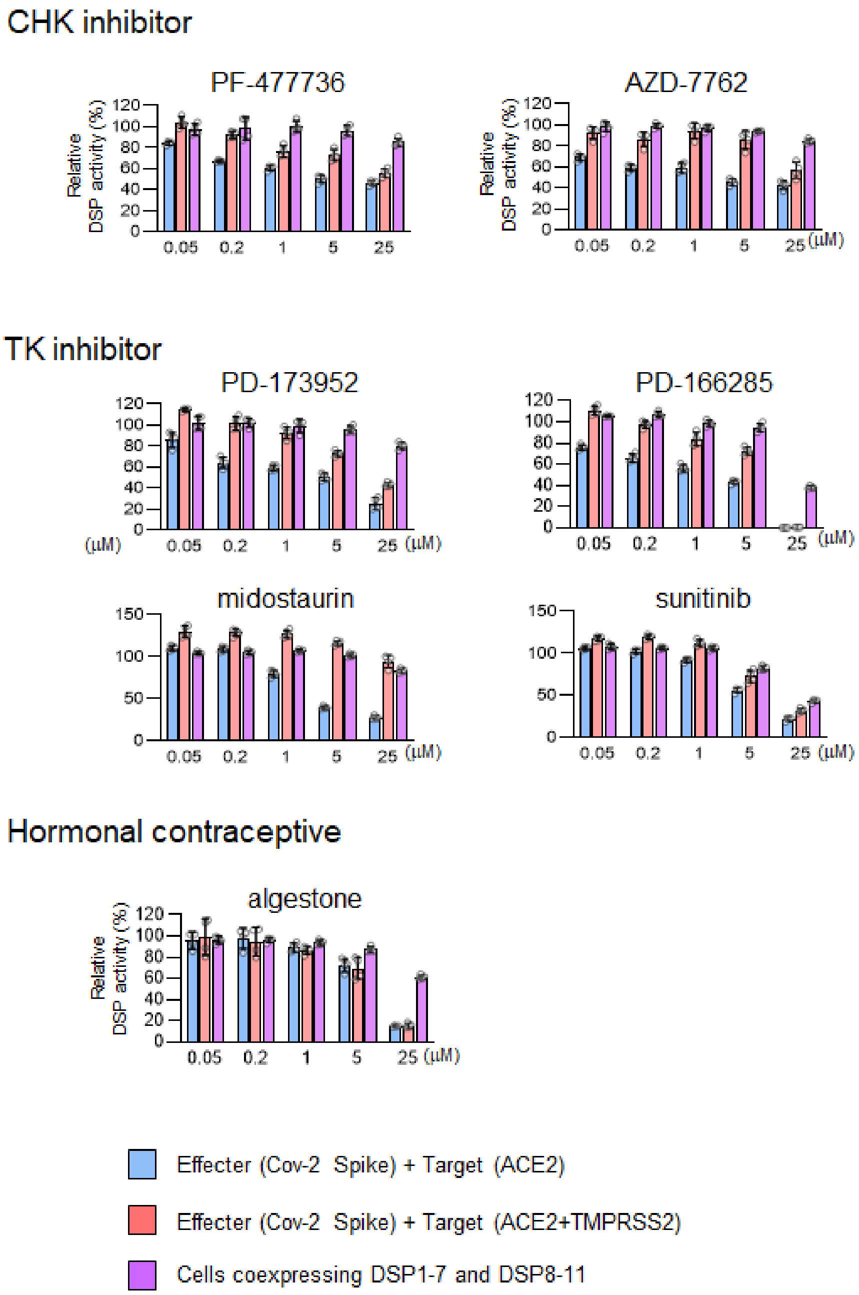


**S3 Fig. Effects of candidate compounds on the cell-cell fusion and RL activities.** Effector cells expressing SARS-CoV-2 S were co-cultured with target cells expressing ACE2 alone for the TMPRSS2-independent cell-cell fusion assay (blue) or cells expressing ACE2 with TMPRSS2 for the TMPRSS2-dependent cell-cell fusion assay (red) in the presence of candidate compounds for 4 h. Cells expressing DSP1-7 and DSP8-11 in the presence of candidate compounds for 4 h to determine whether compounds directly inhibit RL activities (purple). Relative DSP activity was calculated by normalizing the RL activity for each condition to that of the control assay (DMSO alone; set to 100%). Values are means ± SD (*n* = 3/group).

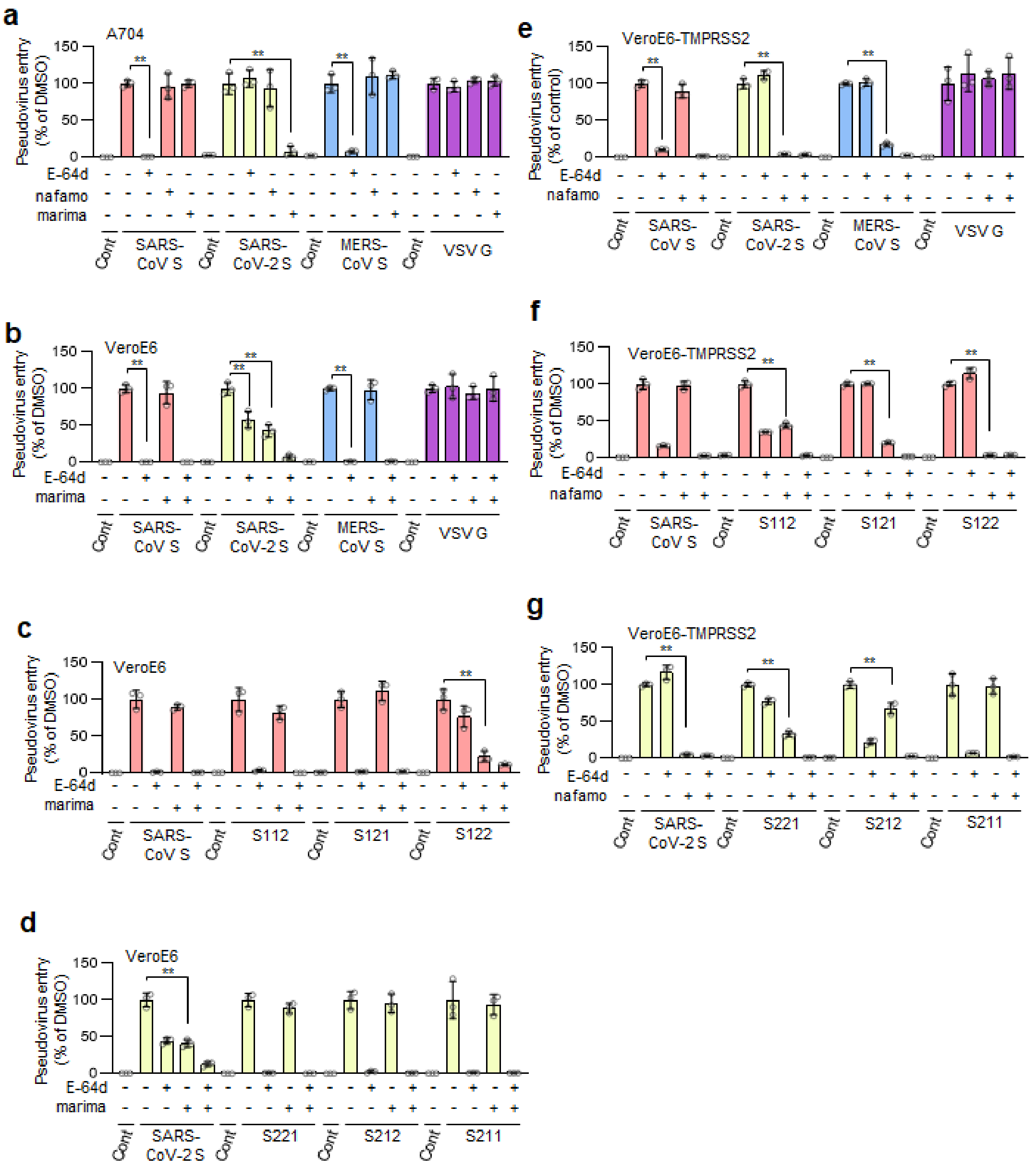


**S4 Fig. The metalloproteinase-dependent entry pathway strictly requires both the furin-cleavage site and S2 region of S protein of SARS-CoV-2.**

Effects of the drugs on the entry of S protein-bearing vesicular stomatitis virus (VSV) pseudotype virus. The relative pseudovirus entry was calculated by normalizing the FL activity for each condition to the FL activity of the cells infected with pseudovirus in the presence of DMSO alone, which was set to 100%. Values are means ± SD (*n* = 3/group). ** $p < 0.01$. Cont: cells infected with pseudovirus without S protein. E-64d: 25 µM E-64d, marima: 1 µM marimastat, nafamo: 10 µM nafamostat. (**a, b**) Effects of E-64d and marimastat on the entry of pseudoviruses bearing SARS-CoV S, SARS-CoV-2 S, MERS-CoV S, or VSV G in A704 (a) and VeroE6 (b) cells. (**c, d**) Effects of E-64d and marimastat on the entry of pseudoviruses bearing chimeric S proteins in VeroE6 cells. (**e**) Effects of E-64d and nafamostat on the entry of pseudoviruses bearing SARS-CoV S, SARS-CoV-2 S, MERS-CoV S, or VSV G in VeroE6-TMPRSS2 cells. (**f, g**) Effects of E-64d and nafamostat on the entry of pseudoviruses bearing chimeric S proteins in VeroE6-TMPRSS2 cells. To establish VeroE6 cells expressing TMPRSS2 (VeroE6-TMPRSS2), recombinant pseudotype lentivirus expressing TMPRSS2 was produced using 293T cells with a VSV-G-expressing plasmid. Cells infected with pseudotype viruses were selected with 300 μg/mL hygromycin for at least 1 week.

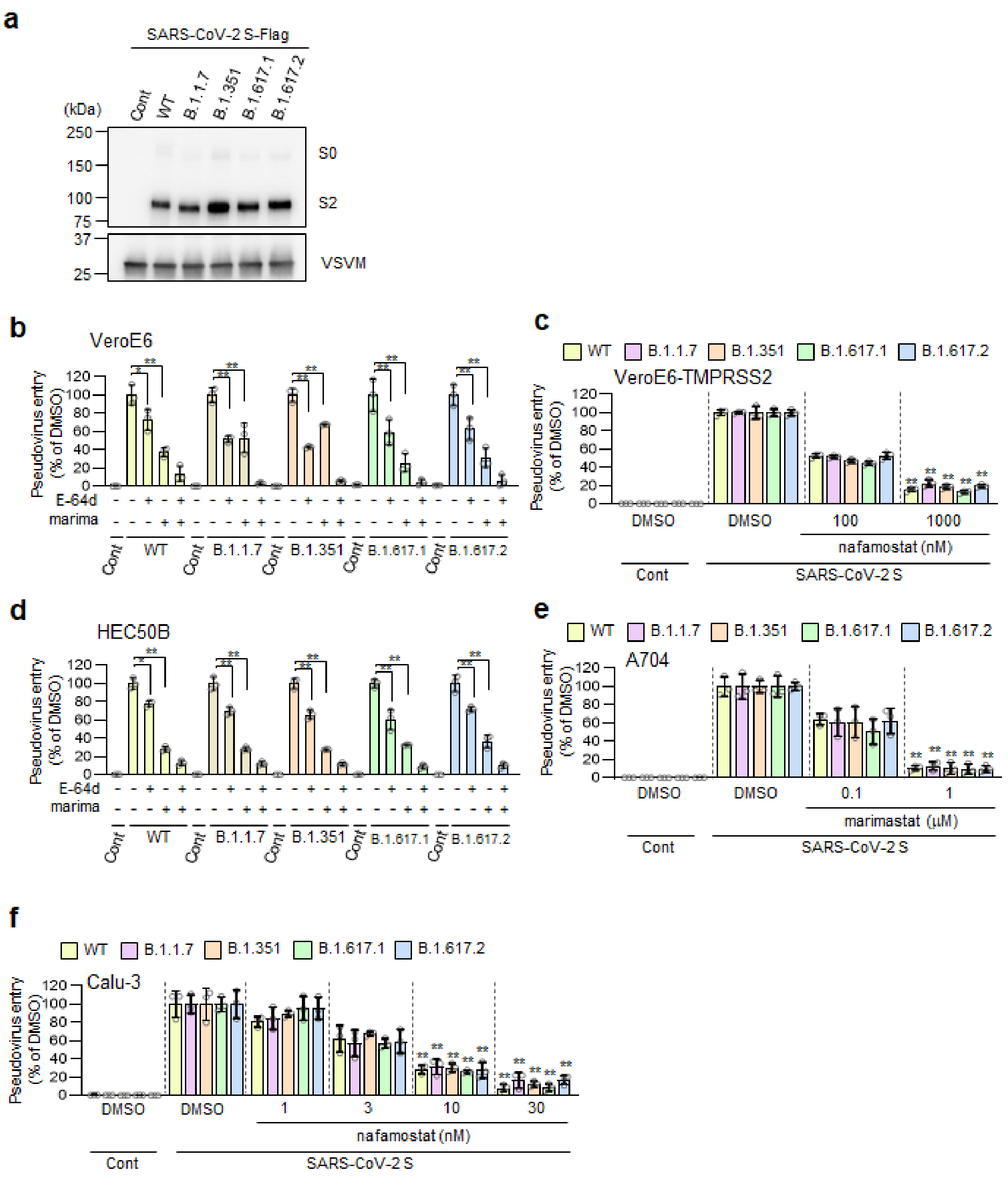


**S5 Fig. Patterns of entry pathways were conserved in various variants of SARS-CoV-2.**

(**a**) Expression of WT or mutant SARS-CoV-2 S proteins with mutations precent in B.1.1.7, B.1.351, B.1.617.1 and B.1.617.2 variants in the pseudoviruses. S proteins were detected using an anti-Flag-tag antibody that binds to a Flag-tag on the C-terminus of S proteins (top). Detection of vesicular stomatitis virus matrix protein (VSV M) served as a control (bottom). S0: uncleaved S protein; S2: cleaved S2 domain of the S protein. (**b**) Effects of E-64d and marimastat on the entry of pseudoviruses bearing SARS-CoV-2 S in VeroE6 cells. E-64d: 25 μM E-64d, marima: 1 μM marimastat. (**c**) Effects of nafamostat on the entry of pseudovirus bearing SARS-CoV-2 S in VeroE6-TMPRSS2 cells. (**d**) Effects of E-64d and marimastat on the entry of pseudoviruses bearing SARS-CoV-2 S in HEC50B cells. E-64d: 25 μM E-64d, marima: 1 μM marimastat. (**e**) Effects of marimastat on the entry of pseudoviruses bearing SARS-CoV-2 S in A704 cells. (**f**) Effects of nafamostat on the entry of pseudoviruses bearing SARS-CoV-2 S in Calu-3 cells. The relative pseudovirus entry was calculated by normalizing the FL activity for each condition to the FL activity of cells infected with pseudovirus in the presence of DMSO alone, which was set to 100%. Values are means ± SD ($n$ = 3/group in b-f). Data were compared with those obtained from cells infected pseudoviruses bearing variant SARS-CoV-2 S in the presence of DMSO alone. * $p < 0.05$, ** $p < 0.01$. Cont: cells infected with a pseudovirus without S protein in (**b-f)**.

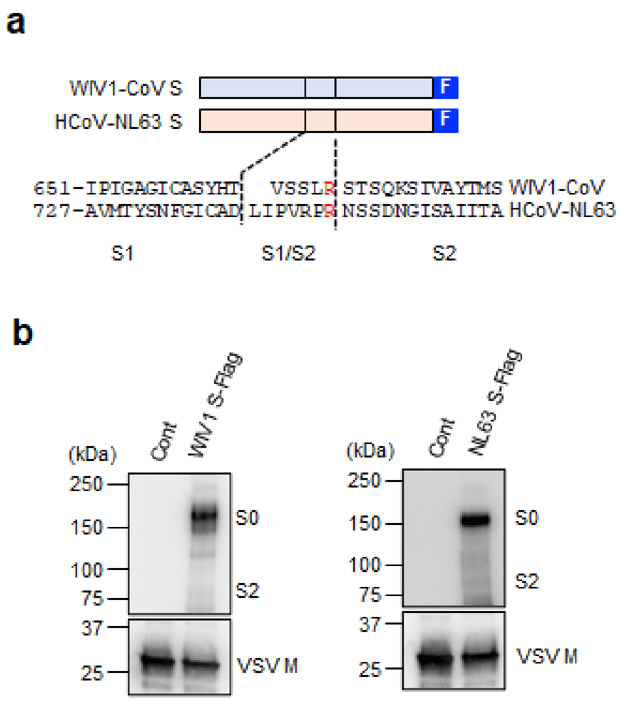


**S6 Fig. Expression of WIV1-CoV and HCoV-NL63 S protein in pseudoviruses.**

(**a**) Schematic illustration of C-terminally FLAG-tagged S proteins of WIV1-CoV and HCoV-NL63 and amino acid sequences of the residues around the S1/S2 boundary of the coronaviruses (bottom). Numbers refer to amino acid residues. F: Flag tag. Arginine residues in the S1/S2 cleavage site and furin cleavage motif are highlighted in red. (**b**) Expression of S protein in pseudoviruses S proteins were detected using an anti-Flag-tag antibody that binds to a Flag-tag on the C-terminus of S proteins (top). The detection of VSV M served as a control (bottom). S0: uncleaved S protein; S2: cleaved S2 domain of the S protein.

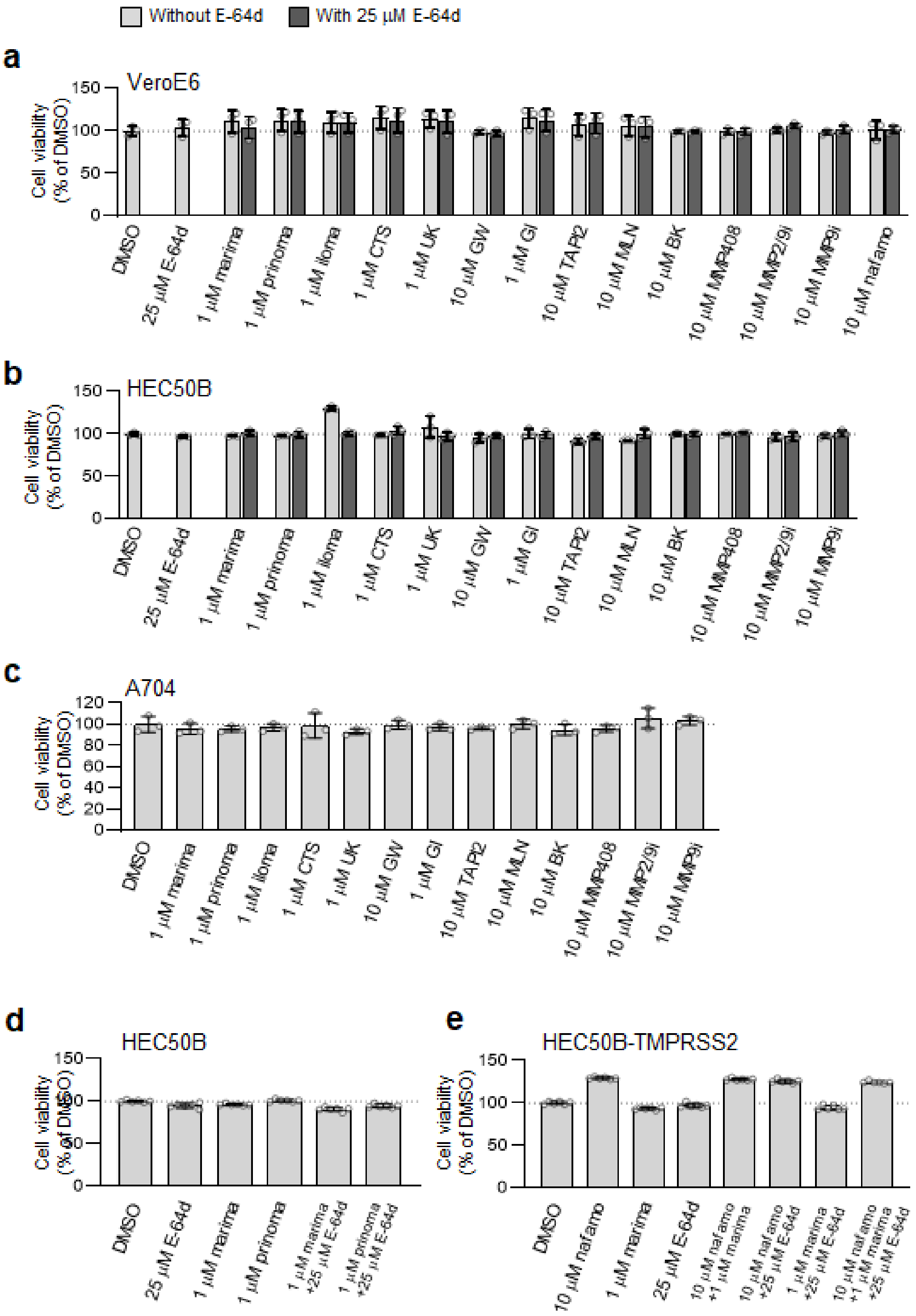


**S7 Fig. Effects of drugs on cell viabilities.**

**(a-c)** VeroE6 (a), HEC50B (b), and A704 (c) cells were treated with various drugs, and cell viability was analyzed using Celltiter-Glo 24 h after the treatment. The relative cell viability was calculated by normalizing the FL activity for each condition to the FL activity of the cells in the presence of DMSO alone, which was set to 100%. Values are means ± SD (*n* = 3/group). (**d, e**) HEC50B (d), and HEC50B-TMPRSS2 (e) cells were treated with various drugs for 3 days. Half of the culture supernatant was replaced daily with fresh medium containing the drugs. The relative cell viability was calculated by normalizing the FL activity for each condition to the FL activity of cells in the presence of DMSO alone, which was set to 100%. Values are means ± SD (*n* = 6/group).

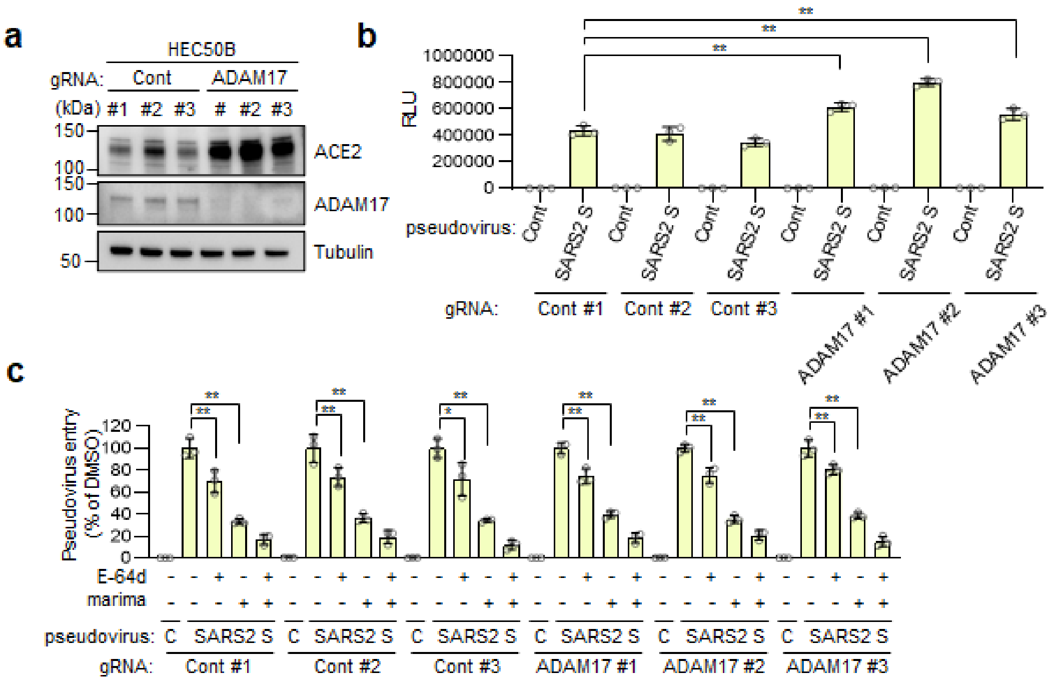


**S8 Fig. Patterns of the entry pathways of the pseudovirus bearing SARS-CoV-2 S were not affected by the ADAM17 knockout in the HEC50B cells.**

(**a**) Effect of the ADAM17 knockout on ACE2 (top), ADAM17 (middle), and tubulin (bottom). (**b**) Effect of the ADAM17 knockout on the entry of the pseudoviruses bearing SARS-CoV-2 S. Values are means ± SD (*n* = 3/group). ** $p < 0.01$. (**c**) Effect of the ADAM17 knockout on the patterns of the entry pathways of SARS-CoV-2 S pseudovirus in HEC50B cells. The relative pseudovirus entry was calculated by normalizing the FL activity for each condition to the FL activity of cells infected with pseudovirus in the presence of DMSO alone, which was set to 100%. Values are means ± SD (*n* = 3/group). * $p < 0.05$, ** $p < 0.01$. E-64d: 25 µM E-64d, marima: 1 µM marimastat. To establish the ADAM17-knockout HEC50B cells, lentiviruses were produced by transfecting the lentiCRISPRv2 vector (#52961 Addgene, MA, USA) with the following gRNA sequences. The gRNA sequences used were 5′-GCG AGG TAT TCG GCT CCG CG-3' (Cont #1), 5′-GCT TTC ACG GAG GTT CGA CG-3' (Cont #2) and 5′-ATG TTG CAG TTC GGC TCG AT-3' (Cont #3) for the control experiments, and 5′-AAC GTT CAG TAC TTG ATG TC-3' (ADAM10 #1) and 5′-GGA CTT CTT CAC TGG ACA CG-3' (ADAM10 #2) and 5′-CTT AAG GTG AGC CTG ACT CT-3' (ADAM10 #3) for the establishment of ADAM17-knockout cells. Pooled HEC50B cells infected with pseudotype viruses were selected with 1 µg/mL puromycin for 1 week.

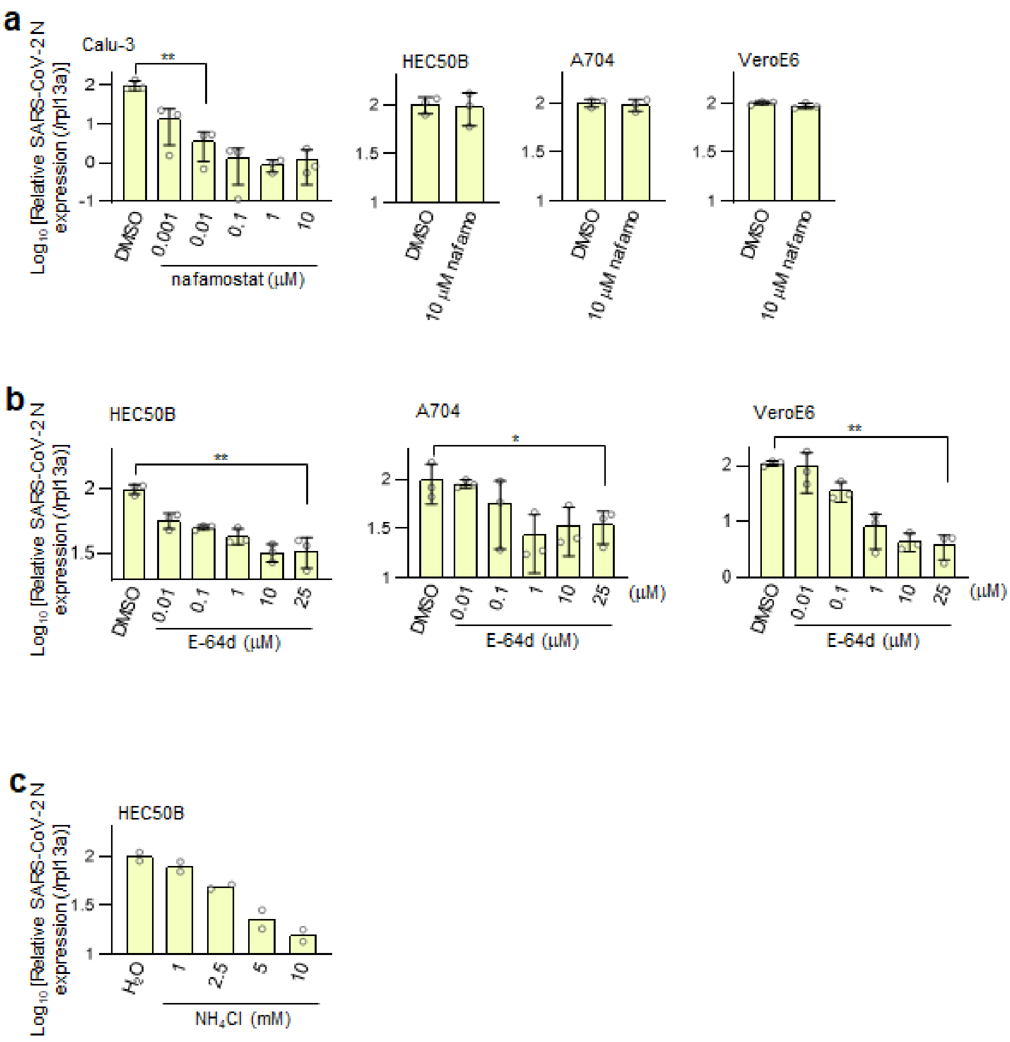


**S9 Fig. Effects of drugs on SARS-CoV-2 infection.**

(**a**) Effects of the nafamostat on the SARS-CoV-2 infection in Calu-3, HEC50B, A704, and VeroE6 cells. Values are means ± SD (*n* = 3/group). ** p < 0.01. (**b**) Effects of the E-64d on the SARS-CoV-2 infection in HEC50B, A704, and VeroE6 cells. Values are means ± SD (*n* = 3/group). * p < 0.05, ** p < 0.01. (**c**) Effects of the $NH_4Cl$ on the SARS-CoV-2 infection in HEC50B cells. Values are means (*n* = 2/group). The relative amount of viral RNA in the cells was normalized to cellular *Rpl13a* mRNA expression in (a-c).

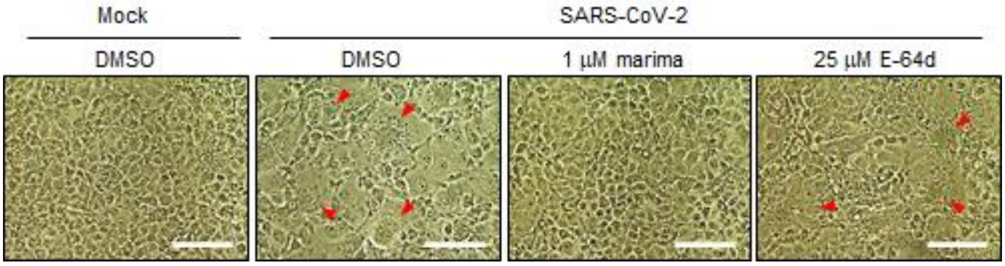


**S10 Fig. The metalloproteinase-dependent entry pathway of authentic SARS-CoV-2 is involved in syncytia formation.**

Phase contrast images of syncytia formation 24 h after SARS-CoV-2 infection in the presence of inhibitors. Red arrowheads indicate syncytia formation Scale bars, 100 μm.
